# TAp73 mediates anti-tumor immunity through regulation of lipid metabolism in the lung tumor microenvironment

**DOI:** 10.1101/2025.11.21.689807

**Authors:** Hayley D. Ackerman, Vanessa Y. Rubio, Andrew J. Davis, John H. Lockhart, Nicole Hackel, Rachel V. Jimenez, Rosa A. Sierra-Mondragon, Jaden R. Baldwin, Christina Carr, Michelle Reiser, Marco Napoli, Xiaoqing Yu, Chia-Ho Cheng, Paul A. Stewart, Suehelay Acevedo-Acevedo, Rahul Checker, Ioannis Grammatikakis, Xiaohua Su, Yaning Wu, Trey Gould, Alexis Bailey, Lary A. Robinson, Eric B. Haura, John M. Koomen, Paulo C. Rodriguez, Elsa R. Flores

**Affiliations:** Department of Molecular Oncology, Moffitt Cancer Center & Research Institute, Tampa, FL; Cancer Biology and Evolution Program, Moffitt Cancer Center & Research Institute, Tampa, FL; Unilever Research and Development, Trumbull, CT; Department of Immunology, Moffitt Cancer Center & Research Institute, Tampa, FL; Department of Biostatistics and Bioinformatics, Moffitt Cancer Center & Research Institute, Tampa, FL; Huntsman Cancer Institute, University of Utah, Salt Lake City, UT; Department of Nutrition and Integrative Physiology, University of Utah, Salt Lake City, UT; Radiation Biology & Health Sciences Division, Bhabha Atomic Research Centre, Trombay, Mumbai-400085, India; Regulatory RNAs and Cancer Section, Genetics Branch, Center for Cancer Research, NCI, NIH, Bethesda, MD; Department of Stem Cell Transplantation and Cellular Therapy, The University of Texas M.D. Anderson Cancer Center, Houston, TX; Division of Human Genetics, Cincinnati Children’s Hospital Medical Center, Cincinnati, OH; Department of Clinical Research, University of Jamestown, Jamestown, ND; Department of Thoracic Oncology, Moffitt Cancer Center & Research Institute, Tampa, FL

## Abstract

While immunotherapy has become the standard of care for lung adenocarcinoma (LUAD) patients without actionable genomic alterations, only a subset of patients benefits from a long-lasting response to immunotherapy. Activation of p53-related signals has emerged as a potential mediator of the lung tumor microenvironment (TME). Given that mutant-p53 interacts with p73 extensively and TAp73-deficient mice develop LUAD, we engineered a mouse model with conditional deletion of *TAp73* to understand the interactions of the p53 family in the TME and in metabolic pathways that impact anti-tumor immunity. We demonstrated that TAp73 exerts a tumor-suppressive role in *Kras^G12D^*-driven LUAD by regulating lipid metabolism in the TME. We identified a TAp73-driven transcriptional signature involving genes in the arachidonic acid metabolism pathway operational in tumor-associated macrophages that favors T-cell activation and thus anti-tumor immunity. Similar transcriptional changes are seen in macrophages from LUAD patients with p53 mutations and in association with response to immunotherapy.

**SIGNIFICANCE:** There is a need to understand how the LUAD TME impacts patient response to immunotherapy. We identified a transcriptional program enacted by TAp73 in tumor alveolar macrophages that supports T-cell activation. Transcriptional and metabolomic data from LUAD patients supports the relevance of this program in response to immune checkpoint inhibition.

## INTRODUCTION

Lung cancer remains the leading cause of cancer-related deaths worldwide. About 85% of cases are non-small cell lung cancer (NSCLC) with lung adenocarcinoma (LUAD) representing the most common subtype of NSCLC. While immune checkpoint inhibitor (ICI) therapy has become the standard of care for NSCLC patients without pharmaceutically targetable alterations, only a subset of patients benefit from a long-lasting response to ICI immunotherapy^1^. Whether p53 mutational status can predict patient response to ICI therapy is an area of intense investigation; however, the results are inconclusive and vary widely depending on the co-occurring driver mutations^2–4^. One way in which mutant-p53 can impact ICI therapy is by rewiring the tumor microenvironment (TME) toward an immunosuppressive state^5^.

While *TP53* is mutated or deleted in about 50% of LUAD cases, its family members *TP63* and *TP73* are not frequently mutated (5% and 3%, respectively)^6^. p73 has several N- and C-terminal isoforms. The major N-terminal isoforms are due to the presence of alternative promoters and give rise to the TAp73 and ΔNp73 isoforms^7^. The TAp73 isoforms contain an N-terminal acidic transactivation domain that is important for the transcriptional regulation of multiple target genes, including some canonical p53 targets, while the shorter ΔNp73 isoforms lack this transactivation domain^8^. LUAD is the most common type of tumor to spontaneously develop in mice that lack the TAp73 isoforms^9^, suggesting that TAp73 plays an important tumor suppressive role in LUAD.

Mutant-p53 can bind to and inhibit TAp73^10^ blocking its ability to transcriptionally activate genes involved in tumor suppressive pathways; therefore, understanding TAp73 function in the context of both wild-type and mutant p53 is important to develop novel therapeutic strategies in LUAD. Previous studies have implicated the p73 isoforms in the inflammatory response^11^ and in macrophage polarization^12^. Given that mutant p53 interacts with p73 extensively and TAp73 deficient mice develop LUAD, we engineered a novel mouse model with conditional deletion of *TAp73* to understand the interactions of the p53 family in the TME, tumor immunity, and metabolic pathways that impact tumor suppression. The ablation of *TAp73* in both the tumor and TME had a significant impact on Kras^G12D^-driven tumor development and progression. Using flow cytometry and single-cell sequencing we observed an expansion of tumor alveolar macrophages with a concomitant depletion of T cells in *TAp73*-null tumors indicating that loss of TAp73 results in a cold TME. We found arachidonic acid metabolism to be a central mediator of the tumor suppressive role of TAp73 in the TME. We determined using chromatin immunoprecipitation and quantitative real time PCR that TAp73 transcriptionally regulates enzymes in the arachidonic acid pathway in tumor alveolar macrophages. Through *in silico* analysis of LUAD single-cell sequencing datasets and targeted metabolomic assessment serum from NSCLC patients we revealed that the TAp73-driven fatty acid metabolism pathway in tumor-associated macrophages is clinically relevant and important for response to ICI therapy.

## RESULTS

### Low survival rates of LUAD correlate with low levels of *TP73* expression

To assess how *TP73* expression affects overall survival of LUAD, we analyzed microarray data from 400 resected LUADs from patients treated at Moffitt Cancer Center^13^. We found that low *TP73* expression has a significant association with worse overall survival outcome (Fig. 1A). When patients were further stratified based on their *TP53* mutation status, we found that the effect of low *TP73* expression on survival only holds true in patients with wild-type *TP53* (Fig. 1B-1C). These data indicate that *TP73* may play a tumor-suppressive role especially in the context of LUADs with wild-type *TP53*.

**Figure 1.**
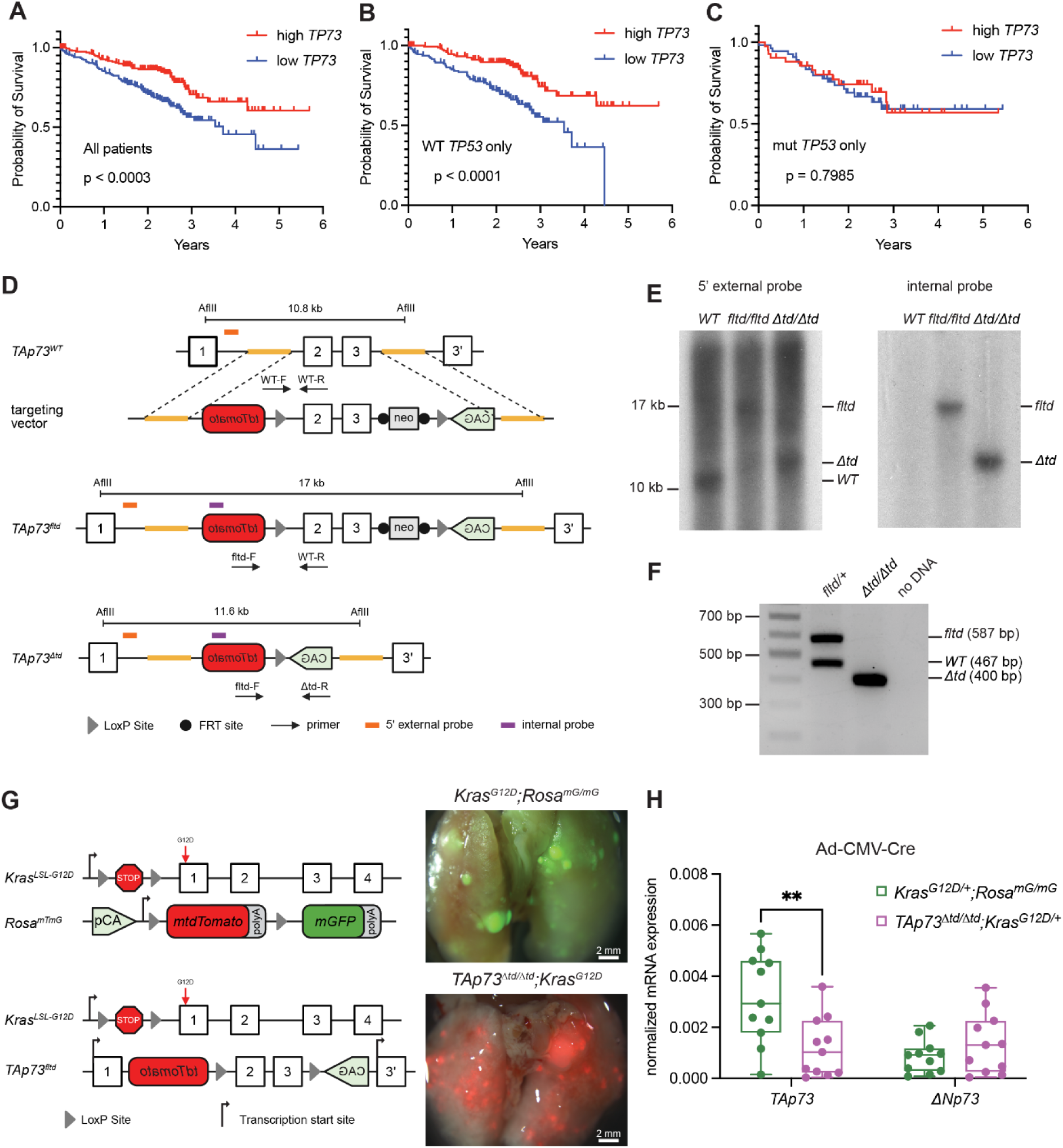
Generation of a *TAp73* conditional knockout (*TAp73^fltd^*) mouse expressing tdTomato in Cre-recombined cells. **A-C,** Kaplan-Meier survival analysis for human LUAD patients (GSE72094) dichotomized by median *TP73* expression. The association between *TP73* expression and overall survival is evaluated among all 398 patients regardless of *TP53* mutational status (**A**), 301 patients with wild-type *TP53* (**B**), and 97 patients with *TP53* mutations (**C**). Statistical comparisons were performed using the log-rank test. **D,** Schematic representation of the targeting vector used to generate the *TAp73^fltd^* allele and the *TAp73^Δtd^* allele that results after Cre-lox recombination. The locations of the Southern blot probes used in **E** and the genotyping primers used in **F** are shown. **E,** Southern blot analysis of genomic DNA from wild-type (WT), *TAp73^fltd/fltd^*, and *TAp73^Δtd/Δtd^* mice using a 5’ external probe (left) and an internal *tdTomato* probe (right). **F,** Genotyping PCR of genomic DNA from *TAp73^fltd/+^* and *TAp73^Δtd/Δtd^* mice. **G,** Schematic representation of the *Kras^LSL-G12D/+^*, *Rosa^mTmG^*, and *TAp73^fltd^* alleles (left) along with representative bright field/fluorescent images of lungs from *Kras^G12D/+^*;*Rosa^mG/mG^* and *TAp73^Δtd/Δtd^*;*Kras^G12D/+^* mice 30 weeks after intratracheal infection with Ad-CMV-Cre (right). Note: the red channel showing non-recombined tissue was omitted from the *Kras^G12D/+^*;*Rosa^mG/mG^* lung for clarity but is shown in Supplementary Fig. S1A. **H,** qPCR of *TAp73* and *ΔNp73* expression relative to *Polr2a* expression in tumors derived from *Kras^G12D/+^*;*Rosa^mG/mG^*and *TAp73^Δtd/Δtd^*;*Kras^G12D/+^* mice infected with Ad-CMV-Cre. \*\**p* < 0.01 by unpaired t test.

### Generation of a *TAp73* conditional knockout (*TAp73^fltd^*) mouse expressing tdTomato in Cre-recombined cells

To understand the role of TAp73 in LUAD, we used the Cre-loxP system to generate a *TAp73* conditional knockout reporter mouse (*TAp73^fltd^*) (Fig. 1D) to enable tissue- and temporal-specific deletion of the *TAp73* isoforms without affecting expression of *ΔNp73* isoforms^14^. LoxP sites were cloned into the endogenous *Trp73* gene flanking exons 2 and 3, which contain the translational start site for the *TAp73* isoforms. The *tdTomato* gene was inserted upstream of the 5’ loxP site in the anti-sense direction, along with a synthetic CAG promoter downstream of the 3’ loxP site so that cells that have undergone Cre-mediated recombination can be identified via tdTomato expression and the resulting red fluorescence. Targeted mouse C57BL/6J embryonic stem cells were screened via Southern blot (Fig. 1E) using probes external to the targeting vector. Properly targeted ES cell clones were injected into donor blastocysts, which were subsequently implanted into pseudopregnant females. The resulting chimeric mice were intercrossed with albino C57BL/6J mice to facilitate the identification of efficient germline transmission, which was subsequently confirmed by Southern blot analysis and PCR (Fig. 1F).

To study the role of *TAp73* in LUAD in the context of wild-type p53, we intercrossed *TAp73^fltd/fltd^* mice with the *Kras^LSL-G12D/+^* model of LUAD which has been shown to recapitulate many of the histological features of human LUAD by 16 weeks after Cre-mediated recombination to activate the Kras^G12D^ mutation^15^ . For *Kras^G12D^* mice without the *TAp73^fltd^* allele, we utilized a conditional fluorescent reporter allele^16^ (*Rosa^mTmG^*) to label and track cells that have undergone Cre-mediated recombination. Non-recombined cells express tdTomato while recombined cells express EGFP (Fig. 1G and Supplementary Fig. S1A-S1B). Adenovirus expressing Cre recombinase was delivered to *Kras^LSL-G12D/+^*;*Rosa^mTmG/mTmG^*and *TAp73^fltd/fltd^*;*Kras^LSL-G12D/+^* mice at 8-10 weeks of age via intratracheal administration, and mice were euthanized 30 weeks later. Quantitative PCR (qPCR) analysis of total RNA extracted from lung tumors collected at endpoint indicates that our targeting strategy ablates *TAp73* expression without affecting the expression of the *ΔNp73* isoforms, a unique feature of this model^9^ (Fig. 1H).

### *TAp73* ablation in the TME has a significant effect on LUAD pathogenesis

To investigate the effects of *TAp73* ablation, we evaluated the lung tumor phenotypes of *Kras^G12D/+^*;*Rosa^mG/mG^*and *TAp73^Δtd/Δtd^*;*Kras^G12D/+^* mice infected with two different adenoviruses expressing Cre recombinase. With Ad-CMV-Cre, Cre recombinase is driven by the ubiquitous CMV promoter which can transduce multiple cell types^17^ allowing us to evaluate the effects of *TAp73* ablation and Kras activation in the TME. With Ad-SPC-Cre, Cre recombinase is specifically expressed in alveolar type II (AT2) cells which are considered the primary cell of origin of LUAD^18^. As alveolar macrophages have previously been shown to be especially affected by Ad-CMV-Cre^17^, we assessed whether they were robustly recombined in our mouse models infected with Ad-CMV-Cre or Ad-SPC-Cre at 30 weeks post-infection. In *Kras^G12D/+^*;*Rosa^mG/mG^*mice (Fig. 2A), the switch from tdTomato to EGFP expression indicated Cre-mediated recombination occurred in Siglec-F^+^ alveolar macrophages (Supplementary Fig. S2A-S2B) isolated from mice infected with Ad-CMV-Cre (Fig. 2B) but not in alveolar macrophages isolated from mice infected with Ad-SPC-Cre (Fig. 2C). Similarly, in *TAp73^Δtd/Δtd^*;*Kras^G12D/+^* mice (Fig. 2D), tdTomato expression indicating Cre-mediated recombination was present in Siglec-F^+^ alveolar macrophages isolated from mice infected with Ad-CMV-Cre (Fig. 2E) but not in alveolar macrophages isolated from mice infected with Ad-SPC-Cre (Fig. 2F). We found that following infection with Ad-CMV-Cre, *TAp73* expression was similarly ablated in both Siglec-F^+^ tumor-associated alveolar macrophages and the remaining Siglec-F^-^ lung cells from tumor-bearing *TAp73^Δtd/Δtd^*;*Kras^G12D/+^*mice (Supplementary Fig. S2C).

**Figure 2.**
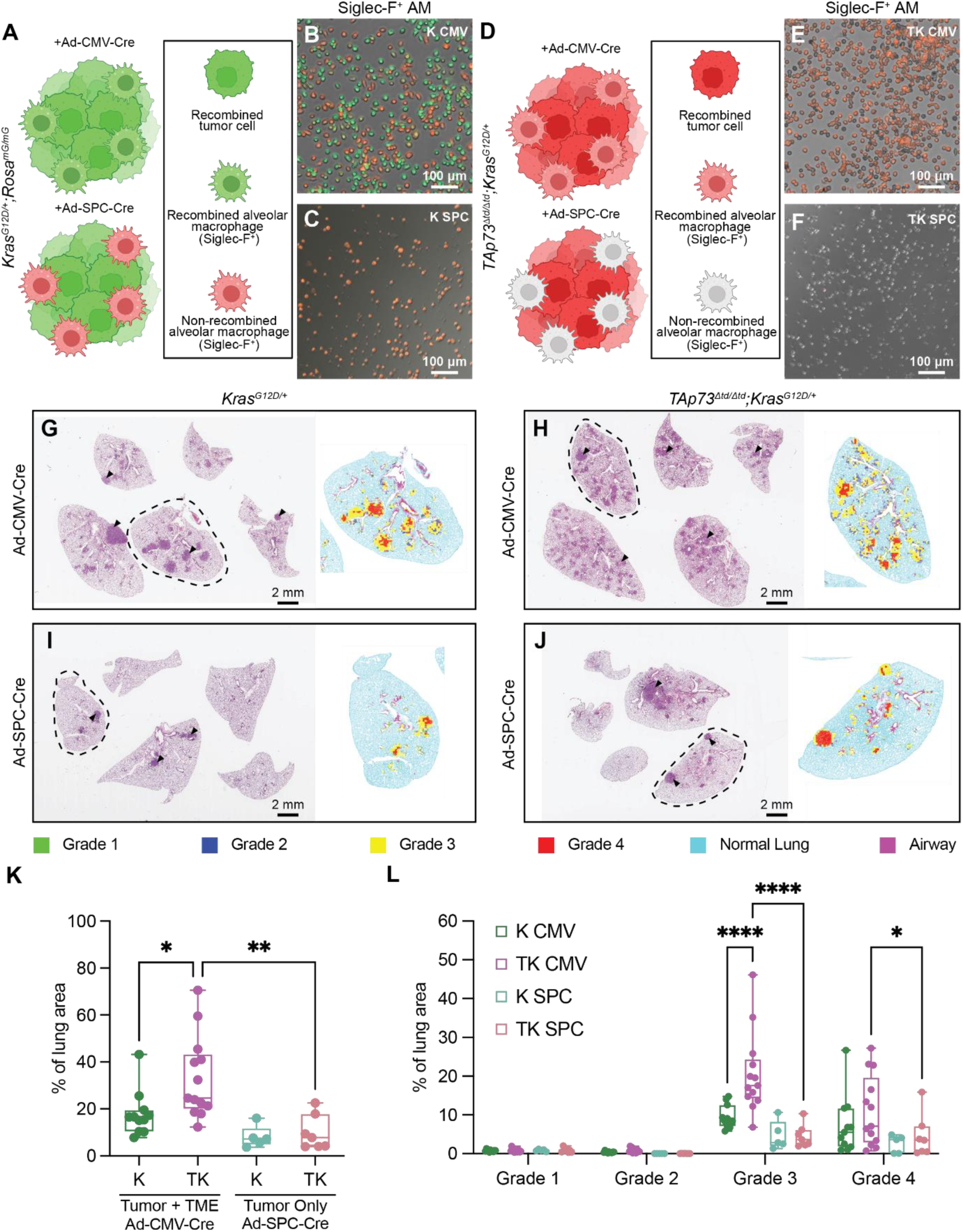
*TAp73* ablation in the TME has a significant effect on LUAD pathogenesis. **A,** Schematic representing the switch from tdTomato to GFP expression in different cell types that are recombined with either Ad-CMV-Cre or Ad-SPC-Cre in *Kras^G12D/+^*;*Rosa^mG/mG^* mice. **B,** Representative fluorescent image of Siglec-F^+^ alveolar macrophages (AM) isolated from a *Kras^G12D/+^*;*Rosa^mG/mG^* mouse infected with Ad-CMV-Cre (K CMV) shows that some cells have been recombined as indicated by the switch from tdTomato to GFP expression. **C,** Representative fluorescent image of Siglec-F^+^ AM isolated from a *Kras^G12D/+^*;*Rosa^mG/mG^* mouse infected with Ad-SPC-Cre (K SPC) shows via the lack of GFP expression that no recombination has occurred. **D,** Schematic representing the induction of tdTomato expression in different cell types that are recombined with either Ad-CMV-Cre or Ad-SPC-Cre in *TAp73^Δtd/Δtd^*;*Kras^G12D/+^*mice. **E,** Representative fluorescent image of Siglec-F^+^ AM isolated from a *TAp73^Δtd/Δtd^*;*Kras^G12D/+^* mouse infected with Ad-CMV-Cre (TK CMV) shows that some cells have been recombined as indicated by the induction of tdTomato expression. **F,** Representative fluorescent image of Siglec-F^+^ AM isolated from a *TAp73^Δtd/Δtd^*;*Kras^G12D/+^*mouse infected with Ad-SPC-Cre (TK SPC) shows via the lack of tdTomato expression that no recombination has occurred. **G-J,** Representative H&E images from *Kras^G12D/+^*;*Rosa^mG/mG^* (**G** and **I**) and *TAp73^Δtd/Δtd^*;*Kras^G12D/+^* (**H** and **J**) mice infected with Ad-CMV-Cre (**G** and **H**) or Ad-SPC-Cre (**I** and **J**). Black arrowheads indicate examples of tumors in each H&E image. Each panel includes the portion of the grading map produced by GLASS-AI for the lobe outlined in the H&E image (see the color key below panels **I-J**). **K** and **L,** Tumor burden (**K**) or tumor burden by grade (**L**) as determined by analysis of H&E images by GLASS-AI. Each tumor was assigned a single grade based on the highest grade present that comprised at least 10% of the tumor’s area. \**p* < 0.05, \*\**p* < 0.01, \*\*\**p* < 0.001, \*\*\*\**p* < 0.0001 by ANOVA followed by Tukey’s multiple comparisons test.

To determine how *TAp73* ablation in different tumor compartments affects LUAD pathogenesis, we used GLASS-AI^19^, a machine learning algorithm, to analyze images of hematoxylin and eosin (H&E) stained cross sections of LUADs from *Kras^G12D/+^*;*Rosa^mG/mG^*(hereafter *Kras^G12D/+^*) and *TAp73^Δtd/Δtd^*;*Kras^G12D/+^*mice infected with Ad-CMV-Cre or Ad-SPC-Cre (Fig. 2G-2J). From this analysis we found that ablation of *TAp73* in the context of oncogenic *Kras* led to a significant increase in tumor burden (Fig. 2K) and tumor grade (Fig. 2L) only when *TAp73* was ablated in both the tumor cells and the TME with Ad-CMV-Cre. These results suggest that TAp73 regulates important tumor suppressive functions in the TME.

### TAp73 regulates metabolism in the tumor microenvironment (TME)

To characterize the modulation of TAp73 in tumor immunity and to determine which metabolic pathways contribute to tumor progression, we performed expression proteomics, untargeted metabolomics, and untargeted lipidomics on LUADs derived from *Kras^G12D/+^* and *TAp73^Δtd/Δtd^*;*Kras^G12D/+^* mice treated with Ad-CMV-Cre to ablate *TAp73* and activate *Kras* in tumor cells and the TME or with Ad-SPC-Cre to ablate *TAp73* and activate *Kras* in alveolar type II cells only. Regardless of the analyte type in the multi-omic experiment, the tumors grouped primarily by the virus that was used and secondarily by genotype when the top 500 features were analyzed (Fig. 3A-3C) indicating a metabolic re-wiring of the TME in the Ad-CMV-cre infected mice.

**Figure 3.**
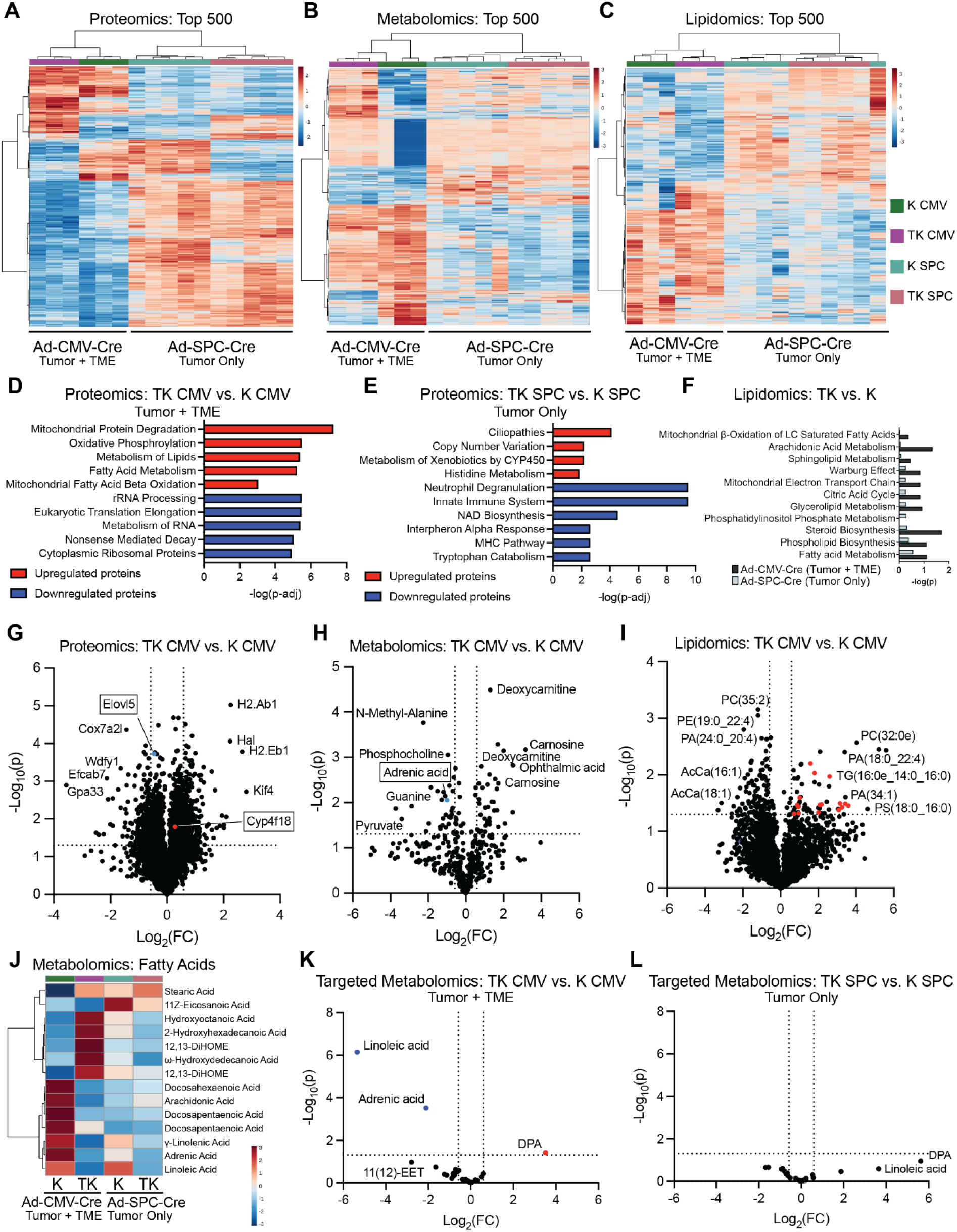
TAp73 regulates metabolism in the TME. **A-C,** Heatmaps representing the relative signal for the top 500 proteins (**A**), metabolites (**B**), or lipids (**C**) that were differentially abundant among tumors isolated from *Kras^G12D/+^* (K) and *TAp73^Δtd/Δtd^*;*Kras^G12D/+^* (TK) mice infected with Ad-CMV-Cre (3 tumors per group) or Ad-SPC-Cre (5 tumors per group). **D** and **E,** Gene set enrichment analysis using mSigDB for proteins that were differentially expressed (p≤0.05, fold change³1.5) between tumors isolated from K and TK mice infected with Ad-CMV-Cre (**D**) or Ad-SPC-Cre (**E**). **F,** Pathway analysis using MetaboAnalyst for differentially abundant lipids between tumors isolated from K and TK mice infected with Ad-CMV-Cre (black) or Ad-SPC-Cre (gray). **G-I,** Volcano plots representing the proteins (**G**), metabolites (**H**), or lipids (**I**) that were differentially abundant among tumors isolated from K and TK mice infected with Ad-CMV-Cre (3 tumors per group). Features were considered to be significantly different if they had a fold change of at least 1.5 with a *p*<0.05. Red dots in **I** represent upregulated lipids in the triglyceride class. **J,** Heatmap representing the average abundance of several fatty acids related to arachidonic acid metabolism that were identified in the untargeted metabolomics dataset represented in panel **B**. **K** and **L,** Volcano plots representing the eicosanoid-targeted metabolomics of tumors isolated from K and TK mice infected with Ad-CMV-Cre (**K**) or Ad-SPC-Cre (**L**), with 6-7 tumors per group. Metabolites were considered to be significantly different if they had a fold change of at least 1.5 with a *p*<0.05.

To understand the tumor suppressive role of TAp73 in the TME, we compared the pathways enriched with *TAp73* ablation in the tumor and TME (Ad-CMV-Cre) with the pathways enriched with tumor-specific/AT2 cell *TAp73* ablation (Ad-SPC-Cre). With the proteomics dataset, pathways involving lipid and fatty acid metabolism were significantly enriched with *TAp73* ablation in the tumor and TME (Fig. 3D) while these pathways were not among the top pathways enriched with tumor-specific *TAp73* ablation (Fig. 3E). With the lipidomics dataset, we found that almost all the pathways were more significantly altered with *TAp73* ablation in both the tumor and TME than with *TAp73* ablation in tumor cells only (Fig. 3F), particularly arachidonic acid metabolism. Arachidonic acid is metabolized into several different classes of bioactive lipid molecules known as eicosanoids that mediate many cellular processes such as inflammation and proliferation^20^.

We found proteins and analytes related to arachidonic acid metabolism among the top hits of the multi-omic datasets. For example, elongation of very long-chain fatty acid protein 5 (ELOVL5) was identified as being downregulated (Fig. 3G, fold change of -1.4) and adrenic acid was shown to be significantly less abundant (Fig. 3H, fold change of -2.0) with *TAp73* ablation in both the tumor and TME. These findings are linked in that ELOVL5 is one of the primary enzymes responsible for generating adrenic acid directly from arachidonic acid^21^. Several triglycerides are significantly more abundant with *TAp73* ablation in both the tumor and TME (Fig. 3I, red dots) which is consistent with the known link between arachidonic acid metabolism and triglyceride synthesis^22^. All these changes are specific to *TAp73* ablation in both the tumor and TME as they are not among the top changes with tumor-specific *TAp73* ablation via Ad-SPC-Cre recombination (Supplementary Fig. S3A-S3C).

While we detected some differentially abundant fatty acids related to arachidonic acid metabolism in the untargeted metabolomics data (Fig. 3J), we developed a targeted mass spectrometry assay to profile 35 different eicosanoids and their precursors that are not robustly detected by untargeted metabolomics or lipidomics^23^. When this assay was applied to extracts from bulk tumors (Supplementary Table S1), a small number of metabolites including adrenic acid were significantly altered when *TAp73* was ablated in the tumor and TME (Ad-CMV-Cre) (Fig. 3K) while none of the targeted metabolites were significantly altered when *TAp73* was only ablated in the tumor cells (Ad-SPC-Cre) (Fig. 3L). Together, these results suggest that the increase in tumor burden with *TAp73* ablation in the tumor and TME may be due to the known role of arachidonic acid metabolism in tumorigenesis^24^.

We performed Matrix-Assisted Laser Desorption Ionization Mass Spectrometry Imaging (MALDI-MSI) to assess the changes in the spatial distribution of metabolites caused by the ablation of *TAp73* in different tumor compartments. We identified tumor regions of interest from H&E-stained frozen sections (Supplementary Fig. S4A) and correlated them with MALDI images acquired from a slide prepared with a serial section (Supplementary Fig. S4B-S4C). We found that arachidonic acid is more abundant in the tumor than in the surrounding lung tissue. In contrast, ablation of *TAp73* in the tumor and TME results in a more diffuse and heterogeneous arachidonic acid signal (Supplementary Fig. S4B). To further investigate this heterogeneity, we performed immunofluorescence for CD45 to distinguish the immune cells of the TME from the epithelial tumor cells (Supplementary Fig. S4D-S4K). This staining revealed that arachidonic acid (Supplementary Fig. S4L-S4O) and adrenic acid (Supplementary Fig. S4P-S4S) are more abundant in the TAp73-deficient immune cells in the TME (Supplementary Fig. S4M and S4Q), indicating a potential role of arachidonic acid in myeloid cells in the higher tumor burden following *TAp73* ablation.

### *TAp73* ablation in the TME drives expansion of macrophages and depletion of T cells

To assess the TME upon *TAp73* ablation, we profiled LUADs from *Kras^G12D/+^* and *TAp73^Δtd/Δtd^*;*Kras^G12D/+^* mice infected with Ad-CMV-Cre of Ad-SPC-Cre for immune markers using immunohistochemical (IHC) staining. We identified a significant reduction in the average number of tumor-infiltrating CD3^+^ T cells (Fig. 4A-E) and CD11b^+^ myeloid cells (Fig. 4F-4J) only when *TAp73* is ablated in both the tumor and the TME. The abundance of alveolar macrophages in tumor-bearing lungs was assessed using flow cytometry to detect co-expression of the cell surface markers Siglec-F and CD11c (Supplementary Fig. S5). We observed up to a three-fold increase in the number of Siglec-F^+^ CD11c^+^ alveolar macrophages present in the lung when *TAp73* was ablated in both the tumor and TME (Fig. 4K-4L) while there was no difference in the abundance of alveolar macrophages with tumor-specific *TAp73* ablation (Fig. 4M-4N).

**Figure 4.**
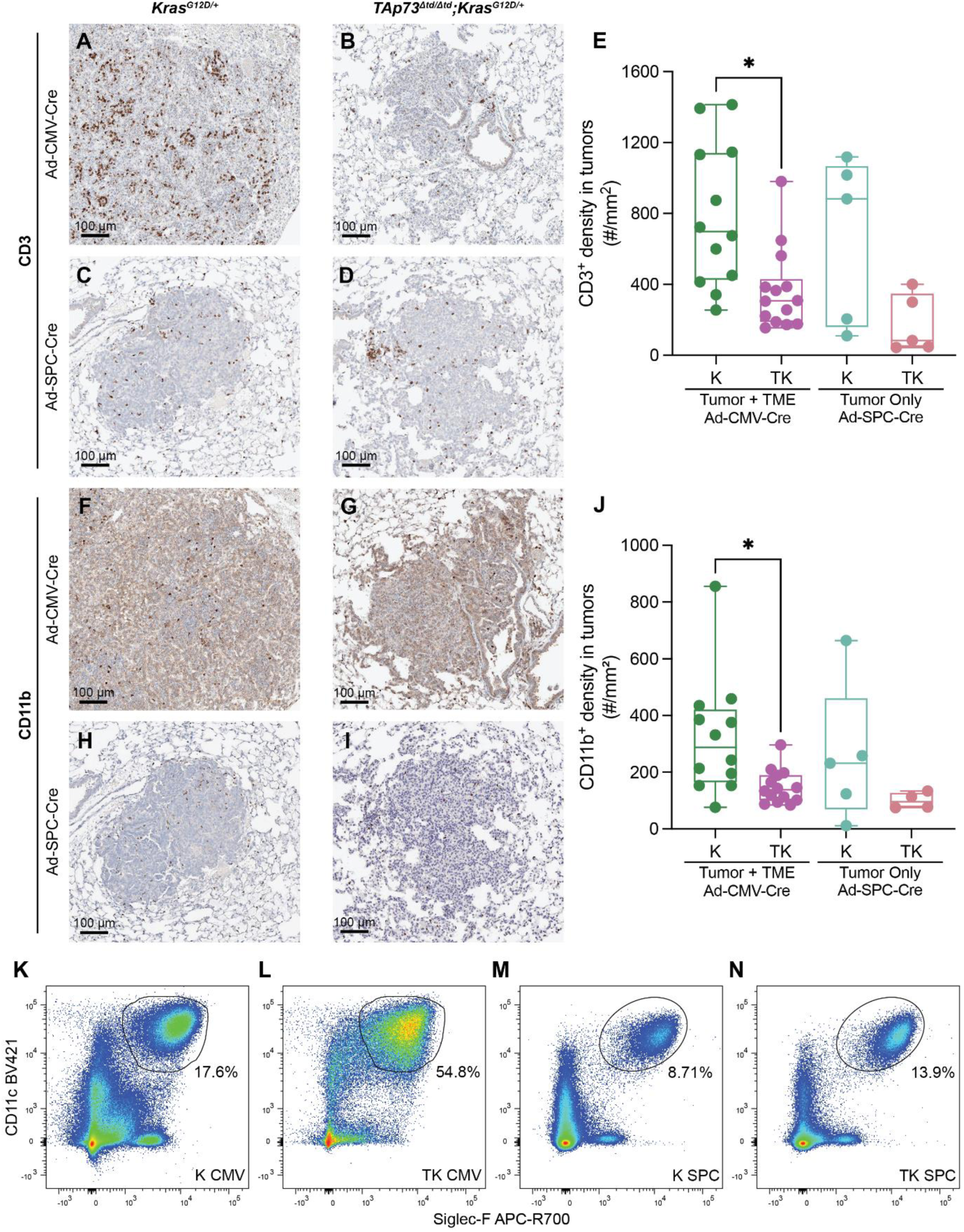
*TAp73* ablation in the TME drives expansion of macrophages and depletion of T cells. **A-D,** Representative CD3 IHC staining in the lungs of *Kras^G12D/+^* (K) and *TAp73^Δtd/Δtd^*;*Kras^G12D/+^* (TK) mice infected with Ad-CMV-Cre (**A** and **B**) or Ad-SPC-Cre (**C** and **D**). **E,** Quantification of CD3 staining represented as the density of CD3^+^ cells within the tumor compartment. \**p* < 0.05 by ANOVA followed by Tukey’s multiple comparisons test. **F-I,** Representative CD11b staining in the lungs of K and TK mice infected with Ad-CMV-Cre (**F** and **G**) or Ad-SPC-Cre (**H** and **I**). **J,** Quantification of CD11b staining represented as the density of CD11b^+^ cells within the tumor compartment. \**p* < 0.05 by ANOVA followed by Tukey’s multiple comparisons test. **K-N,** Representative flow cytometry scatter plots of Siglec-F^+^ CD11c^+^ alveolar macrophages in the lungs of tumor-bearing K (**K** and **M**) or TK (**L** and **N**) mice infected with Ad-CMV-Cre (**K** and **L**) or Ad-SPC-Cre (**M** and **N**). The percentage is calculated based on the CD45^+^ parent population of live single cells.

### Single-cell sequencing identifies an immunosuppressive state in tumor-bearing *TAp73^Δtd/Δtd^*;*Kras^G12D/+^* mice

To further evaluate the cell types of the TME affected by *TAp73* ablation, we performed single-cell sequencing on *Kras^G12D/+^* and *TAp73^Δtd/Δtd^*;*Kras^G12D/+^* mice infected with Ad-CMV-Cre or Ad-SPC-Cre (n=4 of each). The integrated data resolved into 38 unique clusters that could be grouped into 12 major cell types (Fig. 5A). The distribution of the samples across these major cell types was broken down by genotype and whether Cre-recombination occurred in the tumor only (Ad-SPC-Cre) or in the tumor and TME (Ad-CMV-Cre) (Fig. 5B and Supplementary Fig. S6A). There was a statistically significant increase in the percentage of macrophages and statistically significant decreases in the percentage of T cells and neutrophils with the loss of *TAp73* in the tumor and TME (Ad-CMV-Cre). There were no statistically significant differences in the cell type distribution when *TAp73* was lost in the tumor cells only (Ad-SPC-Cre). Macrophage expansion and T cell depletion was also seen in a bulk RNA sequencing dataset from LUADs derived from *Kras^G12D/+^*and *TAp73^Δtd/Δtd^*;*Kras^G12D/+^* mice infected with Ad-CMV-Cre based on GSEA with the cell type signatures identified by single-cell sequencing (Supplementary Fig. S6B). Combined with our immunohistochemical and flow cytometric analyses, this finding confirms that *TAp73* ablation in the tumor and TME leads to an accumulation of macrophages and a depletion of T cells indicative of an immunosuppressive state in LUADs from *TAp73^Δtd/Δtd^*;*Kras^G12D/+^* mice infected with Ad-CMV-Cre.

**Figure 5.**
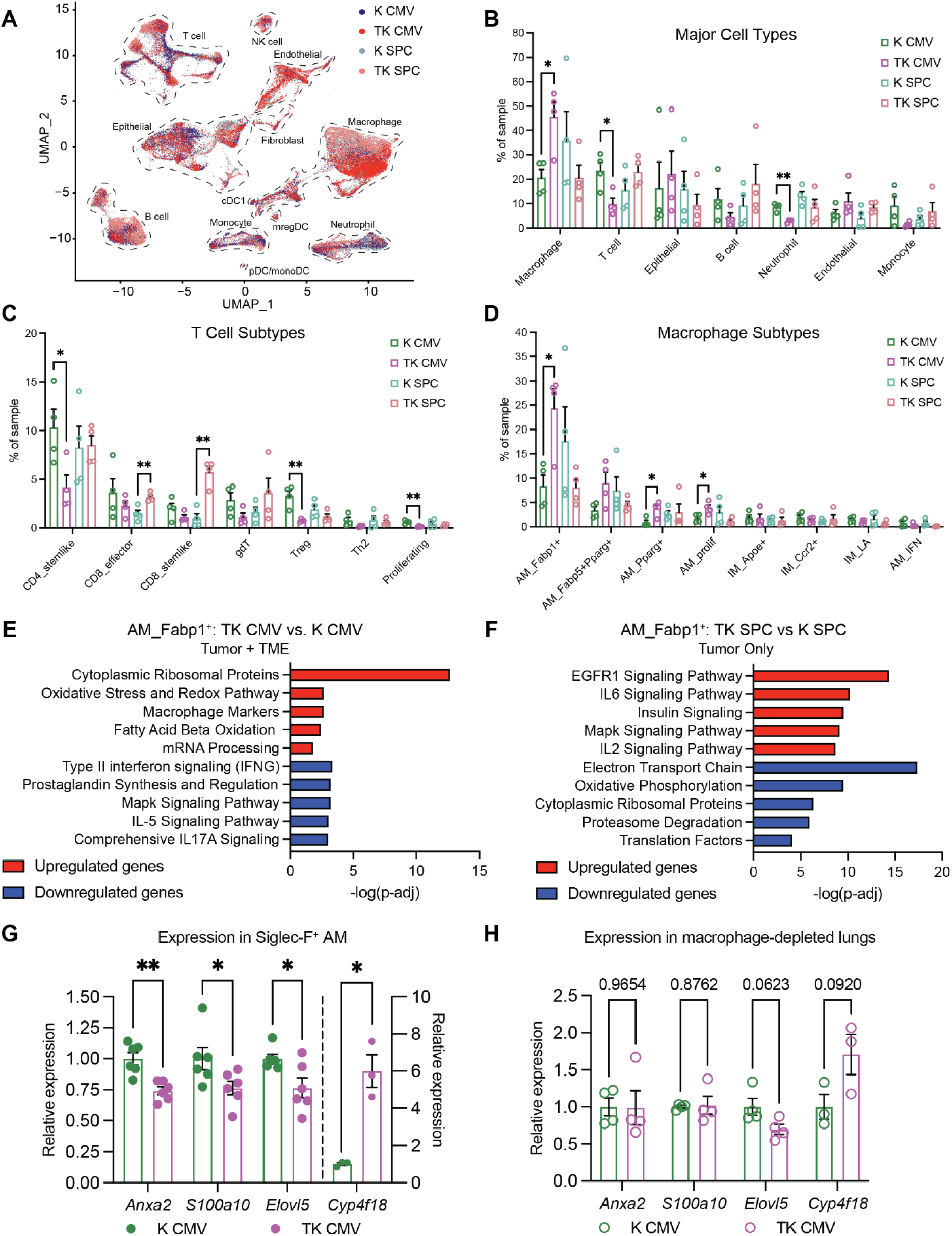
Single-cell sequencing identifies an immunosuppressive state in tumor-bearing *TAp73^Δtd/Δtd^*;*Kras^G12D/+^* mice. **A,** Uniform Manifold Approximation and Projection (UMAP) based on the transcriptional profiles of 75,989 cells derived from 16 samples: 4 samples per genotype:virus combination as color-coded. The cells resolved into 38 clusters which were further grouped into 12 major cell types. **B,** The distribution of the 7 most abundant cell types within each sample across the four genotype:virus combinations. Data are represented as mean ± SEM. The quantification of the remaining 5 cell types is shown in Supplementary Fig. S6A. **C,** The distribution of the 7 different subtypes of T cells within each sample across the four genotype:virus combinations. Data are represented as mean ± SEM. **D,** The distribution of the 8 different subtypes of macrophages within each sample across the four genotype:virus combinations. Data are represented as mean ± SEM. **E** and **F,** Gene set enrichment analysis using the WikiPathways database (mouse, 2024) for genes that were differentially expressed (p-adj≤0.05, fold change > 1.2) between Fabp1^+^ alveolar macrophages from *Kras^G12D/+^*(K) and *TAp73^Δtd/Δtd^*;*Kras^G12D/+^* (TK) mice infected with Ad-CMV-Cre (**E**) or Ad-SPC-Cre (**F**). **G** and **H,** qPCR showing the relative expression of TAp73-target genes in Siglec-F^+^ alveolar macrophages isolated from K and TK mice infected with Ad-CMV-Cre (**G**) and in the remaining macrophage-depleted lung cells (**H**). Data are represented as mean ± SEM. \**p* < 0.05, \*\**p* < 0.01 by multiple unpaired t tests (**B**, **C**, **D,** and **G**).

Further subclustering was performed on the T cells and macrophages (Supplementary Fig. S6C-S6D). There was a significant decrease in the percentage of stemlike CD4^+^ T cells, regulatory T cells (Treg), and proliferating T cells with *TAp73* ablation in the tumor and TME while there was a significant increase in the percentage of effector and stemlike CD8^+^ T cells with tumor-specific *TAp73* ablation (Fig. 5C). Within the macrophages (Fig. 5D), there was a statistically significant increase in the percentage of the predominant alveolar macrophage subtype marked by expression of fatty acid binding protein 1 (Fabp1^+^) as well as significant increases in the Pparg^+^ subtype (peroxisome proliferator-activated receptor gamma) and in the proliferating alveolar macrophages with *TAp73* ablation in the tumor and TME (Ad-CMV-Cre). In contrast, no statistically significant differences were observed in macrophage subtype distribution with the tumor-specific ablation of *TAp73* (Ad-SPC-Cre). Again, these findings indicate that loss of *TAp73* in the TME affects immune cell compositions that may be critical for anti-tumor immunity.

### TAp73-deficient macrophages drive changes in pathways related to arachidonic acid metabolism

To understand the function of the Fabp1^+^ alveolar macrophages, we performed GSEA and generated a heatmap to determine which hallmark gene sets were significantly enriched in the subtypes of alveolar macrophages (Supplementary Fig. S6E). The Fabp1^+^ alveolar macrophages had the highest positive normalized enrichment score (NES) for several pathways including fatty acid metabolism and oxidative phosphorylation, and they were unique in being the only subtype of alveolar macrophage to have a negative NES for the allograft rejection pathway. Additionally, the prostaglandin synthesis pathway involving the conversion of arachidonic acid into bioactive lipid mediators was significantly enriched in the genes that were downregulated in Fabp1^+^ alveolar macrophages when *TAp73* was ablated in both the tumor and TME (Fig. 5E) but not with tumor-specific *TAp73* ablation (Fig. 5F).

To test if TAp73 is transcriptionally regulating genes involved in arachidonic acid metabolism in alveolar macrophages, we performed qPCR on total RNA extracted from Siglec-F^+^ macrophages isolated from tumor-bearing mice. We found that some of the genes in the prostaglandin synthesis pathway identified from single-cell sequencing (Fig. 5E), *Anxa2* and *S100a10*, and other related genes identified from our proteomics dataset (Fig. 3G), *Elovl5* and *Cyp4f18*, were significantly altered in TAp73-deficient alveolar macrophages (Fig. 5G) while they were not significantly altered in the remaining lung cell suspensions from which the macrophages were isolated (Fig. 5H). *Anxa2*, *S100a10*, and *Elovl5* were downregulated in TAp73-deficient alveolar macrophages suggesting they are transcriptionally activated by TAp73. *Cyp4f18* was significantly upregulated in TAp73-depleted macrophages suggesting it is transcriptionally repressed by TAp73.

### TAp73-deficient alveolar macrophages strongly suppress T cells via alterations in the arachidonic acid pathway

The fact that the pathways that were identified as being altered in bulk tumors upon ablation of *TAp73* in the tumor and TME (Fig. 3D and 3F), e.g. arachidonic acid metabolism, overlapped with the pathways identified in TAp73-deficient alveolar macrophages (Fig. 5E) led us to repeat our eicosanoid-targeted metabolomics with Siglec-F^+^ macrophages isolated from tumor-bearing *Kras^G12D/+^* and *TAp73^Δtd/Δtd^*;*Kras^G12D/+^* mice infected with Ad-CMV-Cre to affect the tumor and TME (Fig. 6A and Supplementary Fig. S7A, Supplementary Table S2). The ablation of *TAp73* in alveolar macrophages led to a significant increase in 20-hydroxyeicosatetraenoic acid (20-HETE), 11,12-Epoxyeicosatrienoic acid (11,12-EET), and dihomo-γ-linoleic acid (DGLA) (Fig. 6A). 11,12-EET and 20-HETE are both generated via the cytochrome P450 pathway of arachidonic acid metabolism, while DGLA is a direct precursor of arachidonic acid. None of the targeted metabolites were significantly different between the corresponding Siglec-F^-^ fractions (Supplementary Fig. S7A) indicating that the metabolic signature is significant in tumor-associated macrophages.

**Figure 6.**
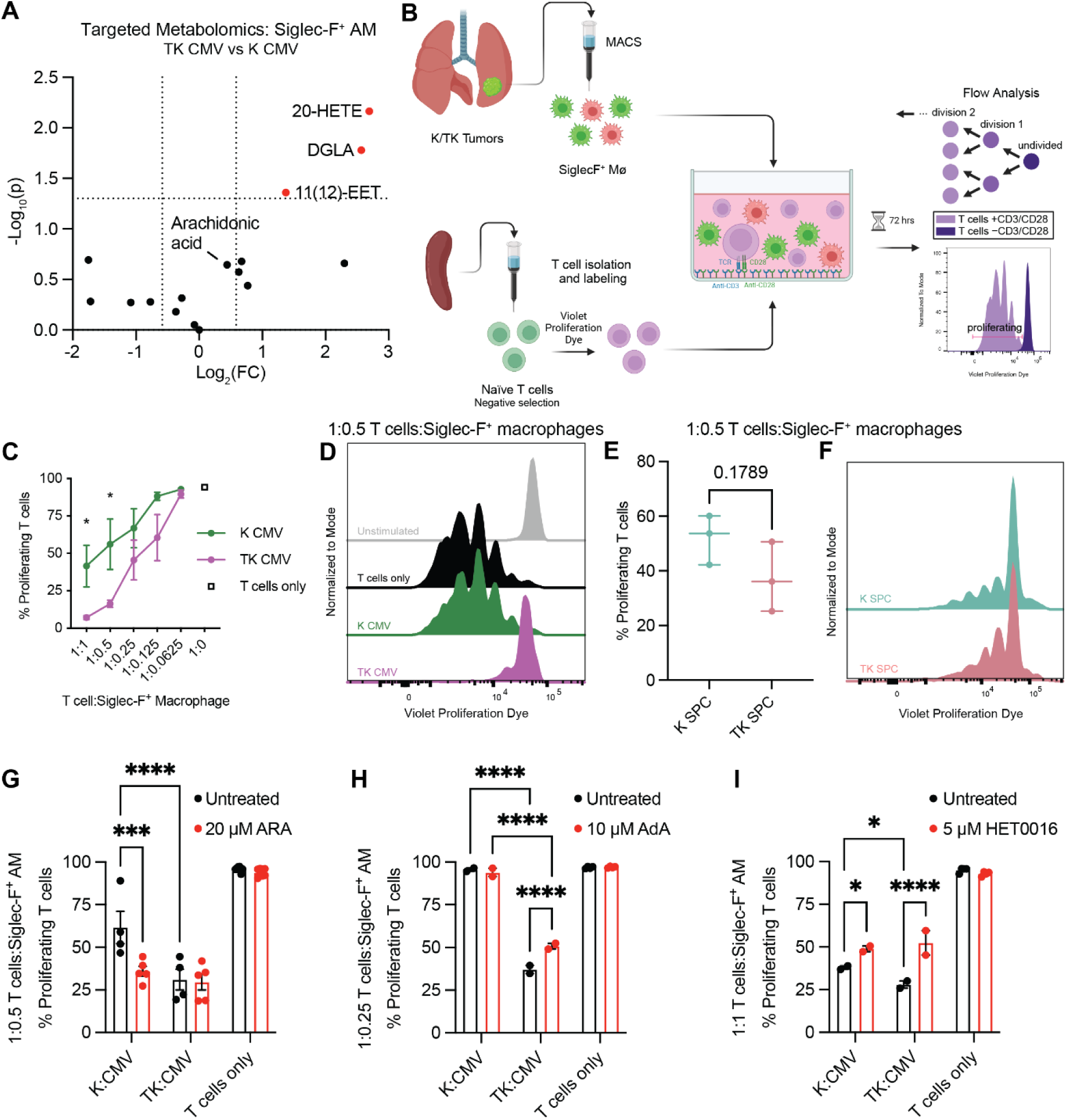
TAp73-deficient macrophages strongly suppress T cells via alterations in the arachidonic acid pathway. **A,** Volcano plot representing the eicosanoid-targeted metabolomics of Siglec-F^+^ alveolar macrophages isolated from *Kras^G12D/+^* (K) and *TAp73^Δtd/Δtd^*;*Kras^G12D/+^* (TK) mice infected with Ad-CMV-Cre. Metabolites were considered to be significantly different if they had a fold change of at least 1.5 with *p*<0.05. **B,** Schematic representation of the assay to test the ability of Siglec-F^+^ alveolar macrophages isolated from tumor-bearing mice to inhibit the proliferation of naïve T cells activated by plate-bound anti-CD3 and anti-CD28 in a three-day co-culture. T cell proliferation was quantified based on the dilution of eFluor450 dye and gated based on T cells cultured alone without CD3/CD28 stimulation. **C,** Proliferation of CD3/CD28-stimulated T cells with serially diluted Siglec-F^+^ alveolar macrophages isolated from K and TK mice infected with Ad-CMV-Cre. Data includes replicates from two independent experiments and are represented as mean ± SEM. **D,** Representative histograms for T cells cultured alone and unstimulated (gray), alone and stimulated with anti-CD3/CD28 (black), or stimulated with anti-CD3/CD28 and cultured at a 1:0.5 ratio of T cells to Siglec-F^+^ macrophages from K (green) or TK (purple) mice infected with Ad-CMV-Cre. **E,** Proliferation of CD3/CD28-stimulated T cells cultured at a 1:0.5 ratio of T cells to Siglec-F^+^ alveolar macrophages isolated from K and TK mice infected with Ad-SPC-Cre. **F,** Representative histograms for the data in **E**. **G-I,** Effect of arachidonic acid (ARA, **G**), adrenic acid (AdA, **H**), or HET0016 (20-HETE inhibitor, **I**) on T cell proliferation when cultured alone or in the presence of Siglec-F^+^ alveolar macrophages isolated from K and TK mice infected with Ad-CMV-Cre. Data are represented as mean ± SEM. \**p* < 0.05, \*\**p* < 0.01, \*\*\**p* < 0.001, \*\*\*\**p* < 0.0001 by multiple unpaired t tests (**C** and **E**) or by ANOVA followed by Fisher’s LSD test (**G**, **H**, and **I**).

To determine the role of TAp73 in creating an immunosuppressive TME, we isolated Siglec-F^+^ alveolar macrophages from *Kras^G12D/+^* and *TAp73^Δtd/Δtd^*;*Kras^G12D/+^* mice infected with Ad-CMV-Cre or Ad-SPC-Cre and assessed their ability to directly suppress wild-type anti-CD3/CD28 T cells in a three-day co-culture of macrophages and T cells (Fig. 6B and Supplementary Fig. S7B). T cell proliferation was quantified as the dilution of a violet cell proliferation dye and gated based on T cells cultured alone without CD3/CD28. Alveolar macrophages from *TAp73^Δtd/Δtd^*;*Kras^G12D/+^* mice infected with Ad-CMV-Cre show a significantly increased suppressive capacity against T cells compared to alveolar macrophages from *Kras^G12D/+^* mice infected with Ad-CMV-Cre when cultured at a 1:1 or 1:0.5 ratio of T cells to macrophages (Fig. 6C-6D). There was not a significant difference in the suppressive capacity of macrophages isolated from *Kras^G12D/+^* and *TAp73^Δtd/Δtd^*;*Kras^G12D/+^* mice infected with Ad-SPC-Cre (Fig. 6E-6F). The inhibition of T cell function by macrophages derived from *TAp73^Δtd/Δtd^*;*Kras^G12D/+^*mice infected with Ad-CMV-Cre is further supported by ELISA data showing that there was a significant decrease in the amount of IFNγ released by the T cells that were co-cultured with macrophages from these mice compared to the T cells co-cultured with macrophages from *Kras^G12D/+^*mice infected with Ad-CMV-Cre (Supplementary Fig. S7C). There was not a significant difference in the levels of IFNγ when the T cells were co-cultured with macrophages isolated from mice infected with Ad-SPC-Cre (Supplementary Fig. S7C). Taken together, these data indicate that alveolar macrophages deficient of TAp73 suppress T cells and in this way may enhance tumorigenesis by creating an immune cold TME.

To investigate the role that altered arachidonic acid metabolism plays in the immunosuppressive phenotype of TAp73-deficient alveolar macrophages, we repeated the T cell suppression assays with supplementation or inhibition of the dysregulated metabolites we identified in this pathway. We found that supplementation with 20 µM arachidonic acid^25^, which is increased after *TAp73* ablation (Fig. 6A), increased the suppressiveness of TAp73-replete macrophages from *Kras^G12D/+^* mice infected with Ad-CMV-Cre in line with the level of suppression of the TAp73-deficient macrophages from *TAp73^Δtd/Δtd^*;*Kras^G12D/+^*mice infected with Ad-CMV-Cre (Fig. 6G and Supplementary Fig. S7D). Supplementation with 10 µM adrenic acid^26^, which is decreased with *TAp73* ablation (Fig. 3K), lowered T cell suppression carried out by TAp73-deficient macrophages from *TAp73^Δtd/Δtd^*;*Kras^G12D/+^*mice infected with Ad-CMV-Cre (Fig. 6H and Supplementary Fig. S7E). Lastly, we found that using 5 µM HET0016^27^ to inhibit the production of 20-HETE, which is increased with *TAp73* ablation (Fig. 6A), decreased T cell suppression induced by macrophages from both *Kras^G12D/+^*and *TAp73^Δtd/Δtd^*;*Kras^G12D/+^* mice infected with Ad-CMV-Cre, though the effect was more robust in macrophages from *TAp73^Δtd/Δtd^*;*Kras^G12D/+^* mice (Fig. 6I and Supplementary Fig. S7F). Notably, none of these treatments affected the proliferation of T cells cultured in the absence of alveolar macrophages (Fig. 6G-6I and Supplementary Fig. S7D-S7F).

### TAp73 transcriptionally regulates enzymes in the arachidonic acid pathway in alveolar macrophages

To determine whether TAp73 directly binds to the promoters of the transcriptional targets identified in Figure 5, we used chromatin immunoprecipitation (ChIP) followed by qPCR on a human macrophage-like cell line (THP-1) engineered to overexpress FLAG-tagged TAp73α (Fig. 7A). We saw that *ANXA2*, *S100A10*, *ELOVL5*, and *CYP4F3* are direct targets of TAp73 (Fig. 7B) as the transcription factor was recruited to predicted p73 binding sites in their promoters or introns. We also assessed the recruitment of TAp73 to a p73 binding site in the third intron of *CYP4F2* as we found that the sequence was highly similar (90%) to the p73 binding site and surrounding DNA in the third intron of *CYP4F3*. We found high levels of TAp73 bound to *CYP4F2* suggesting that its tumor suppressive function may involve the regulation of multiple members of the cytochrome P450 family, especially those with similar function (i.e. production of 20-HETE^28^). We next performed the eicosanoid targeted metabolomics assay on the engineered THP-1 cells both before and after differentiation from monocyte to macrophage with phorbol 12-myristate 13-acetate (PMA) (Fig. 7C, Supplementary Table S3). Overall, levels of arachidonic acid and related metabolites were much higher in cells following differentiation to macrophages, suggesting the relevance of this pathway in the cell type. Within the differentiated cells, we observed that overexpression of TAp73 led to a decrease in arachidonic acid and an increase in adrenic acid in alignment with our proposed model (Fig. 7D). We also saw the concomitant decrease in several prostaglandins that would indicate a less immunosuppressive state with increased *TAp73* expression. Taken together, our data shows that, in alveolar macrophages, TAp73 transcriptionally regulates enzymes in the arachidonic acid pathway to maintain the resulting bioactive lipid mediators in a state that favors T cell activation and thus anti-tumor immunity.

**Figure 7.**
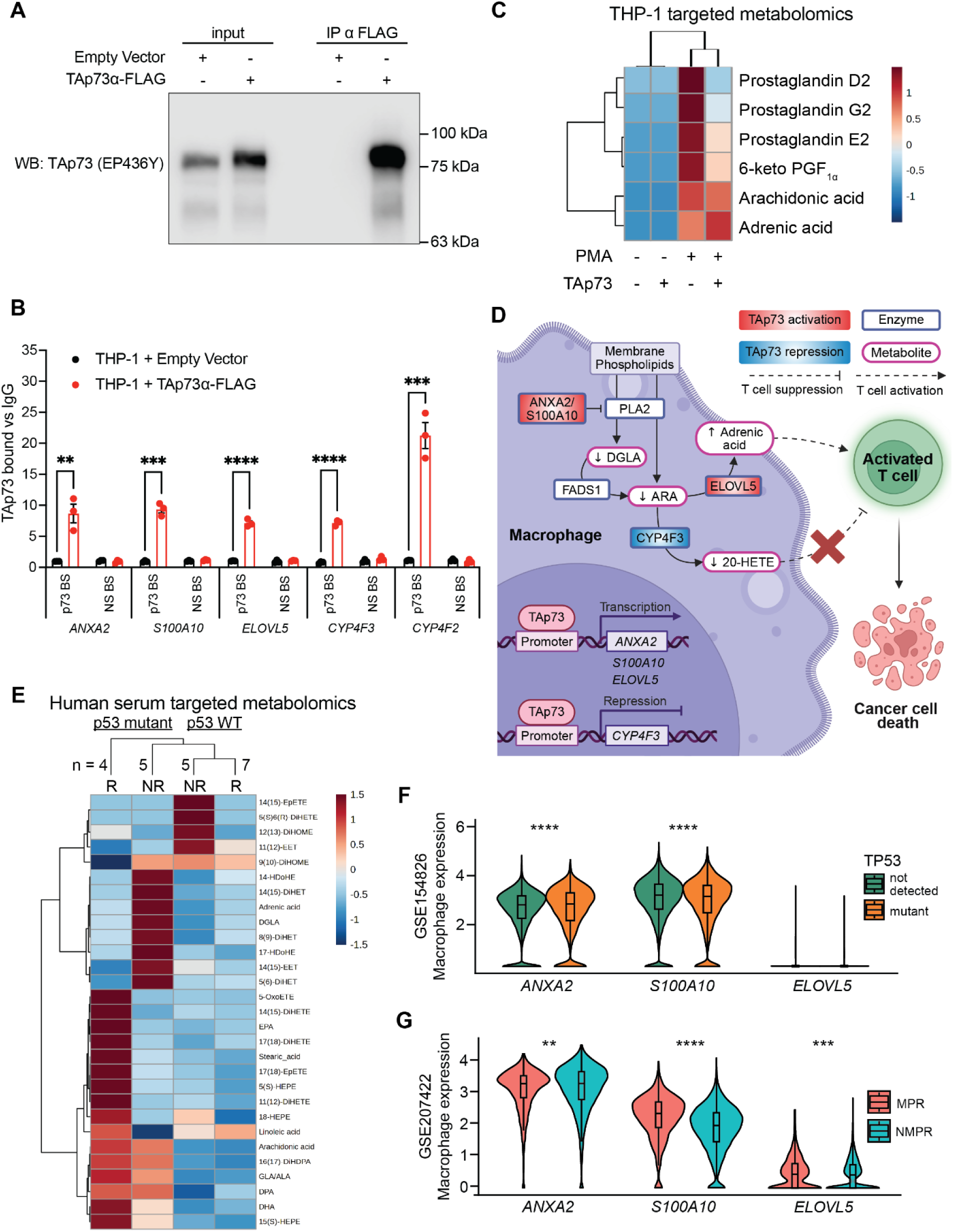
TAp73 transcriptionally regulates enzymes in the arachidonic acid pathway in alveolar macrophages. **A,** Representative Western blot showing the overexpression of TAp73α FLAG inhuman THP-1 cells, as well as the enrichment for TAp73 following immunoprecipitation (IP) with anti-FLAG beads. **B,** qPCR of TAp73 ChIP assays comparing the relative enrichment for p73 binding sites (BS) pulled down by anti-FLAG beads in a TAp73α FLAG overexpression model compared to the enrichment of a non-specific (NS) binding site. Data are represented as mean ± SEM of three independent experiments. **C,** Heatmap representing the average abundance of targeted eicosanoid metabolites detected in THP-1 cells in their monocyte-like state (- PMA) or after differentiation to a macrophage-like state (+ PMA). Cells were transduced with an empty vector (-TAp73) or with TAp73α FLAG (+TAp73). **D,** Working model showing that in alveolar macrophages, TAp73 transcriptionally regulates enzymes in the arachidonic acid pathway to maintain the resulting bioactive lipid mediators in a state that favors T cell activation and thus anti-tumor immunity. PLA2: phospholipase A2, FADS1: fatty acid desaturase 1, DGLA: dihomo-γ-linoleic acid, ARA: arachidonic acid, 20-HETE: 20-hydroxyeicosatetraenoic acid **E,** Heatmap representing the average abundance of targeted eicosanoid metabolites in serum isolated from human LUAD patients (MCC 18611). The patients were grouped based on their p53 mutational status as well as by those who responded (R) or did not respond (NR) to immune checkpoint inhibitor therapy. The number of patients per group is indicated above the heatmap. **F,** Violin plots showing expression levels of TAp73 target genes in macrophages from different LUAD patients according to their p53 mutational status (GSE154826). Data is from 10 patients with p53 mutations and 20 patients where p53 mutations were not detected. **G,** Violin plots showing expression levels of TAp73 target genes in macrophages from different LUAD patients according to their response to immunotherapy (GSE207422). Data is from 2 patients with a major pathologic response (MPR) to immunotherapy and 4 patients with a non-major pathologic response (NMPR). MPR is defined as having no more than 10% residual viable tumor cells by H&E staining after therapy. \**p* < 0.05, \*\**p* < 0.01, \*\*\**p* < 0.001, \*\*\*\**p* < 0.0001 by multiple unpaired t tests (**B**) or by Kruskal-Wallis test (**F** and **G**).

### Metabolic signature in serum from p53-mutant LUAD patients treated with immunotherapy matches TAp73-regulated pathways

To determine the clinical relevance of the TAp73-driven signature in tumor alveolar macrophages that drive anti-tumor immunity, we used samples derived from patients treated with immunotherapy. Previous transcriptomic analysis of tumors from NSCLC patients showed that the fatty acid metabolism pathway (including the *S100A10* and *ELOVL5* genes) was enriched in patients who responded to ICI therapy while patients whose tumors were resistant to therapy had a strong association with a tumor-associated macrophage gene signature^29^. This observation led us to interrogate a panel of 21 serum samples from human NSCLC patients^30^ with the eicosanoid-targeted metabolomics protocol to determine if the changes identified from our macrophage-rich *TAp73^Δtd/Δtd^*;*Kras^G12D/+^*mice infected with Ad-CMV-Cre with a poor anti-tumor immune response may correspond to changes seen between patients who either did or did not respond to ICI therapy (Fig. 7E, Supplementary Table S4). Patients who responded either had complete response, partial response, or stable disease for at least 12 months based on RECIST guidelines^31^. We further stratified the patients based on their p53 mutational status as p53 mutations are known to inhibit TAp73^10^ and thus patients with p53 mutations would more closely resemble the TAp73-low phenotype we characterized in mice. Overall, we found that the arachidonic acid pathway was much more active in patients with p53 mutations. Among p53-mutant patients, we saw that a decrease in DGLA was associated with response as we would have predicted based on our results in macrophages derived from *TAp73^Δtd/Δtd^*;*Kras^G12D/+^* mice infected with Ad-CMV-Cre (Fig. 6A and 7D).

To specifically identify a role for our TAp73 signature in tumor-associated macrophages from human LUADs, we interrogated publicly available single-cell sequencing datasets from LUAD patients^32–34^. We found that *S100A10* was significantly decreased in macrophages from 10 patients with p53 mutations as compared to 20 patients with wild-type p53^32^ (Fig. 7F). Both *S100A10* and *ELOVL5* were significantly increased in macrophages from 2 patients who responded to ICI therapy as compared to 4 patients who failed to respond^33^ (Fig. 7G). In another dataset looking at different nodules within a single patient treated with immunotherapy^34^, we found that the fatty acid metabolism signature that was reported to be enriched in the nodule that responded to immunotherapy was driven by expression in cells of the monocyte/macrophage lineage (Supplementary Fig. S8A, analyzed on the TISCH2 platform^35^). In this dataset, *ANXA2*, *S100A10*, and *ELOVL5* were differentially expressed in the macrophages from the different nodules with *ANXA2* being significantly increased in the nodule that responded to immunotherapy (Supplementary Fig. S8B). In summary, our data show that the arachidonic acid pathway driven by TAp73 is altered in tumor associated macrophages from patients with p53 mutations and is relevant for response to ICI therapy.

## DISCUSSION

While it is widely appreciated that the TME plays a critical role in determining patient outcomes^36,37^, technical challenges have limited studies of the interaction between a tumor and its microenvironment *in vivo*. Here, we demonstrate that TAp73 exerts a tumor suppressive role in *Kras^G12D^*-driven LUAD by regulating lipid metabolism in the TME and specifically in tumor-associated macrophages. By administering Cre-recombinase using adenoviral vectors with distinct tissue and cell-type specificity, we discovered that *TAp73* ablation in both the tumor and TME led to a significant increase in tumor burden and grade, while the effects of tumor cell-specific *TAp73* ablation were mild suggesting a crucial role for the tumor suppressive activities of TAp73 in the TME. Further, we combined multi-omic analyses of bulk tumors, MALDI-MSI spatial metabolomics, and single-cell RNA sequencing to determine that the alveolar macrophages were driving the TAp73-regulated changes in the TME. These changes centered on the arachidonic acid metabolism pathway, with TAp73 directly regulating the transcription of enzymes that affect levels of bioactive lipid mediators (e.g., adrenic acid and 20-HETE). Additionally, ablation of *TAp73* in the tumor and TME led to an expansion of alveolar macrophages and a depletion of T cells – two findings that appear to be linked via the highly immunosuppressive nature of the TAp73-deficient macrophages that we unveiled. Supplementing the TAp73-deficient macrophages with adrenic acid or inhibiting their production of 20-HETE rescued the immunosuppression, thus cementing the importance of the arachidonic acid pathway in the overall tumor-promoting phenotype. Lastly, we found that the TAp73-driven fatty acid metabolism pathway in tumor-associated macrophages impacts response to immune checkpoint therapy.

We hypothesized that TAp73 plays an important role in the TME as previous studies have implicated the p73 isoforms in the inflammatory response^11^, and TAp73 in particular is thought to play a role in regulating macrophage polarization^12^. However, until now there has not been a model that has allowed for temporal and spatial control of *TAp73* ablation. With our Cre-dependent knockout allele we avoid any of the developmental defects that could confound the results in a constitutive *TAp73* knockout model. We can also control the cell and tissue specificity depending on which adeno-Cre virus is used. This selectivity is important as we see in our data that TAp73 drives distinct transcriptional programs in different cell types, as is known to be the case for its family member p53^38^. While *TAp73* is not frequently mutated in LUAD, about 30% of TP53 missense mutations are gain-of-function mutations capable of inhibiting the activity of TAp73^39^. This likely explains why in our analysis of a human microarray dataset we see that TAp73 expression only correlates with survival in patients with wild-type p53, as those with p53 mutations may have diminished TAp73 function regardless of its transcriptional levels. While the mechanism is not yet well established, recent literature suggests that exosomes released from tumor cells harboring p53 mutations can influence the pro-tumor functions of the TME^40^. There is even some data indicating that the mutated p53 protein itself is part of the exosomal cargo and can be taken up by cells in the TME^41^, thus supporting the relevance to understanding TAp73 function in tumor-associated macrophages.

We show that in tumor-associated alveolar macrophages, TAp73 activates the expression of *ANXA2*, *S100A10*, and *ELOVL5* and represses the transcription of *CYP4F3* (the human ortholog of mouse *Cyp4f18*). At normal levels, ANXA2 and S100A10 form a heterotetramer that can prevent phospholipase A2 from releasing arachidonic acid and/or DGLA from the cellular membrane^42^. Downstream of arachidonic acid, increased levels of adrenic acid via ELOVL5 and decreased levels of 20-HETE via CYP4F3 promotes a state of T cell activation as shown by our T cell:macrophage co-culture where the supplementation or inhibition of these metabolites decreased the T cell suppression carried out by TAp73-deficient macrophages. There is an abundance of literature describing the utility of the HET0016 compound as an inhibitor of the CYP4 family of enzymes with decreased 20-HETE levels correlating with decreased cancer cell proliferation, angiogenesis, and/or metastasis in various cancer subtypes^28^. More recently, a mechanism has been proposed by which CYP4F2-driven 20-HETE levels in NSCLC led to immune evasion by inducing PD-L1 expression in cancer-associated fibroblasts^27^. Our work supports this mechanism, though our findings identify alveolar macrophages as the cell type responsible for mediating the immunosuppression of T cells via their production of 20-HETE and ultimately influencing tumor development. It has proven difficult to characterize the many different enzymes of the cytochrome p450 family as the family is expanded in mice compared to humans^28^, but *Cyp4f18* has been characterized as an alveolar macrophage signature gene^43^. The p73 binding site and surrounding sequence in the third intron of *CYP4F3* is almost identical to the sequence of the third intron of *CYP4F2*. As we demonstrate that TAp73 is significantly recruited to both of these binding sites, it implies that the transcriptional control of these functionally related enzymes by TAp73 has been conserved.

ICI therapy is invaluable for treating the many NSCLC patients without pharmacologically targetable driver mutations, but only a minority of patients have a long-lasting response to immunotherapy^1^. Recently, studies have shown that the composition of the TME may be a valuable predictor of response to ICI therapy^32,44^. For example, one study found that a signature for CD8^+^ exhausted T cells was strongly associated with response to therapy while a signature for monocytes/macrophages was strongly associated with resistance to therapy^29^. These trends are consistent with our mouse models where we found that the macrophage expansion and T cell depletion that occurs after *TAp73* ablation in the tumor and TME correlates with a more aggressive tumor phenotype. Response to ICI therapy has also been linked to fatty acid metabolism^29^, with the TAp73 targets *ELOVL5* and *S100A10* identified in our study among the genes driving the enrichment of this pathway in responders. However, with bulk transcriptomic data there is no way to know which cell types were driving this pathway. To overcome this limitation, we performed *in silico* analysis of single-cell sequencing datasets from LUAD patients^32–34^. From this analysis, we found that fatty acid metabolism is primarily driven by macrophages. It also revealed that the TAp73-targets *ANXA2*, *S100A10, and ELOVL5* are differentially expressed in macrophages in the TME of LUADs depending on the p53 mutational status of the patient’s tumor and in association with the patient’s response to ICI therapy.

Interestingly, in a cohort of NSCLC patients treated with ICI therapy we saw that eicosanoid levels in the serum were much higher in patients with tumors harboring p53 mutations. This metabolic data supports our proposed mechanism in which the inhibition of TAp73 transcriptional activity by mutant-p53 leads to increased activity of the arachidonic acid pathway in macrophages and impacts anti-tumor immunity. Overall, our findings suggest that recent immunoengineering methods involving macrophage-based cell therapies^45,46^ could be leveraged to alter arachidonic acid metabolism in the TME in order to overcome the immune evasion driven by mutant-p53^5^ and improve the efficacy of immune checkpoint inhibitor therapy in LUAD.

## METHODS

### Human datasets

Gene expression array data from resected tumors from 442 LUAD patients (GEO GSE72094) as well as clinical data of the patients (survival, stage, annotation of prevalent driver mutations and tumor suppressor mutations, etc.) were kindly provided by Dr. Eric Haura^13^. CEL files were normalized against their median sample using IRON^47^. An RNA-quality related batch-effect was corrected by a Partial Least Squares (PLS) model. The log-rank test was used to evaluate the association of gene expression dichotomized by median with overall survival based on a p-value < 0.05.

Serum samples from LUAD patients were kindly provided by Dr. Lary Robinson. The samples were collected from a prospective observational cohort study, which involved patients treated for metastatic NSCLC at H. Lee Moffitt Cancer Center and Research Institute^30^. Adult patients were enrolled with histologically confirmed, newly diagnosed metastatic NSCLC who were scheduled to undergo immune checkpoint inhibitor therapy with or without chemotherapy. Informed consent was obtained from all patients, and the study was approved by the Advarra Institutional Review Board (MCC 18611, PRO00017235). We limited our cohort to serum samples from patients that had been evaluated for p53 mutational status.

### Animal studies

Mouse studies were approved by the IACUC at the University of Texas MD Anderson Cancer Center or at the Moffitt Cancer Center (IS00013710). The following mice were purchased from The Jackson Laboratory: C57BL/6J (RRID:IMSR_JAX:000664), *Kras^LSL-G12D^* (B6.129S4-*Kras^tm4Tyj^*/J, RRID:IMSR_JAX:008179), and *Rosa^mTmG/mTmG^* (B6.129(Cg)-*Gt(ROSA)26Sor^tm4(ACTB-tdTomato,-EGFP)Luo^*/J, RRID:IMSR_JAX:007676). Age- and sex-matched mice were used for all experiments.

### Generating *TAp73^fltd/fltd^* mice

The cre-loxP strategy was used to generate the TAp73 conditional knockout reporter allele (*TAp73^fltd^*). Genomic DNA encompassing the *Trp73* locus from intron 1 to intron 3 was amplified from mouse genomic DNA (C57BL/6). A neomycin resistance gene (neo) flanked by frt sites was inserted in intron 3. LoxP sites were cloned into the endogenous locus upstream (5’) to exon 2 and downstream (3’) of the frt-flanked neo cassette. *tdTomato* was cloned upstream of the 5’ loxP site in an antisense orientation, while the synthetic CAG promoter was cloned, in an antisense orientation, downstream of the 3’ loxP site. The modified *p73* locus was cloned into the pL253 plasmid vector^48^. Mouse embryonic stem cells (G4) electroporated with the targeting vector were analyzed by Southern blot for proper targeting of the *TAp73^fltd^* allele. Resulting chimeras were mated with C57BL/6 albino females and genotyped as described below. Mice with germline transmission of the targeted allele (*TAp73^fltd^*) were intercrossed to generate homozygous mice (TAp73^fltd/fltd^).

### Intratracheal adenoviral infections

Intratracheal administration of viral particles was carried out as described previously^49^. Briefly, 8-10 week-old mice were anesthetized via an intraperitoneal injection of 100 mg/kg ketamine with 10 mg/kg xylazine. They were then intratracheally infected with 7.5E7 PFU per mouse of adenovirus expressing Cre precipitated with 10 mM CaCl_2_ in a final volume of 50 µl minimum essential media (MEM, Sigma M4655). Ad5-CMV-Cre was purchased from Baylor College of Medicine while Ad5-SPC-Cre was purchased from the University of Iowa. Following delivery of Ad-Cre, mice were given a subcutaneous injection of 1 mg/kg of atipamezole in 0.9% NaCl to reverse the sedative effects. Mice were then allowed to recover on a heating pad and monitored until consciousness was fully regained.

### Cell lines

The human monocytic leukemia cell line THP-1 was purchased from ATCC (TIB-202, RRID:CVCL_0006). The cells were maintained in suspension in a humidified incubator at 37 °C with 5% carbon dioxide and subcultured under sterile conditions according to the instructions from ATCC. Cells were cultured in complete RPMI 1640 (GenClone 25-506) supplemented with 10% heat-inactivated fetal bovine serum (Sigma-Aldrich F0926), 1% penicillin/streptomycin (Gibco 15-140-122), and 0.05 mM 2-mercaptoethanol (Sigma-Aldrich M3148). The cells were authenticated by short tandem repeat profiling and tested once a month to ensure they were negative for mycoplasma contamination.

We purchased two custom plasmids from System Biosciences: an empty vector control plasmid (pCDH-MSCV-MCS-EF1α-RFP-Puro) and a plasmid to stably overexpress human TAp73α (pCDH-MSCV-TAp73α-EF1α-RFP-Puro). We used PCR extension and restriction cloning to add a FLAG tag to the C-terminus of TAp73α. The generated constructs (3 μg) were transfected into 293T cells along with 1 μg of each vector required for lentivirus packaging (pCMV-VSVG and psPAX2) using X-tremeGENE HP DNA Transfection Reagent (Roche 6366244001) according to the manufacturer’s protocol. Supernatant containing lentivirus was collected from the transfected cells after 48 hours, filtered (0.45 µm), and added to target THP-1 cells for 24 hours in the presence of 10 μg/ml Polybrene (Santa Cruz sc-134220). After infection, cells were selected for at least 7 days with 1 μg/ml puromycin (Sigma-Aldrich P8833).

### Southern blot

Genomic DNA was digested overnight with AflII and separated on a 0.8% agarose gel. After denaturation, it was transferred and crosslinked to a nylon membrane for hybridization with radiolabeled probes (Sequences in Table S5).

### Genotyping PCR

Genomic DNA isolated from tail snips or from cells was analyzed by PCR using the following primers all at 0.5 μM: TAp73WT-F (5’-GGGTACTTGTCCTGTGTCATCCTC-3’), TAp73WT-R (5’-AACCGCTTTCCCCACGATTTCCTC-3’), TAp73fltd-F (5’-GGGATGTCGGCGGGGTGCTTCA-3’), and TAp73Δtd-R (5’-CGTGCTGGTTATTGTGCTGTCTCA-3’).

### Microscopy

Bright field and fluorescent images of whole lungs were taken on a Zeiss Axio Zoom.V16 microscope while images of isolated macrophages were taken on a Zeiss Axio Observer.Z1. **qPCR**

Total RNA was isolated from snap frozen tumors or from isolated cells using the miRNeasy Mini Kit (Qiagen 217004). cDNA was synthesized using qScript Ultra SuperMix (Quantabio 95217-500) according to the manufacturer’s protocol. qPCR reactions were performed in triplicate using the QuantStudio 6 Flex real-time PCR system with TaqMan Universal PCR Master Mix and TaqMan Gene Expression Assays (Applied Biosystems). The following TaqMan assays were used: Polr2a (Mm00839502_m1), TAp73 (Mm00660223_m1), ΔNp73 (Mm01263882_m1), Anxa2 (Mm01150673_m1), Elovl5 (Mm00506717_m1), S100a10 (Mm00501458_g1), and Cyp4f18 (Mm00499348_m1).

### Alveolar macrophage isolation

Tumor-bearing mouse lungs were collected into PBS with 2% FBS and kept on ice until ready for processing. To generate a single-cell suspension, the tissue was first minced with a razor blade and then digested for 30 minutes at 37° with 1 mg/mL Collagenase IV (Worthington LS004188) and 25 U/mL DNase I (Sigma D4527) diluted in HBSS (Corning 21-023-CV). The digestion media was neutralized by 3- to 4-fold dilution with PBS containing 2% fetal bovine serum (FBS) and filtered through 70 µm mesh. The cells were washed with 5 mL PBS containing 2% FBS and 1 mM EDTA and counted before being subjected to immunomagnetic positive selection for Siglec-F^+^ cells. The Siglec-F^+^ cells were enriched by using the EasySep Release Mouse APC-Positive Selection Kit (StemCell 100-0033) according to the manufacturer’s protocol along with 1 µg Siglec-F APC antibody (Miltenyi 130-123-816, RRID:AB_2811558) per mL of sample.

### Flow cytometry

A single-cell suspension was generated from tumor-bearing mouse lungs as described above. After filtration, the red blood cells were lysed (Biolegend 420301) and the remaining cells were washed with cold PBS. Cells were then stained with a near-IR viability dye (Invitrogen L34975) according to the manufacturer’s protocol. Cells were then treated with TruStain FcX PLUS (BioLegend 156604) and stained on ice for 30 minutes in the dark with the following antibody cocktail diluted in BD Horizon Brilliant Stain Buffer (BD 563794): anti-CD45 BUV395 (BD 565967, RRID:AB_2651134), anti-CD11c BV421 (BD 565451, RRID:AB_2744278), anti Siglec-F APC (Miltenyi 130-123-816, RRID:AB_2811558). Cells were then washed with FACS buffer (PBS + 1% BSA) and transferred to flow tubes through a 35 µm strainer cap. Data was acquired on a BD LSRII and analyzed with FlowJo version 10 according to the gating strategy in Supplementary Fig. S5.

Co-cultured macrophages and T cells were stained with anti-CD45 APC (BioLegend 103112, RRID:AB_312977) and TruStain FcX (BioLegend 101320, RRID: AB_1574975). Data was acquired on a Beckman Coulter CytoFlex II and analyzed with FlowJo version 11 according to the gating strategy in Supplementary Fig. S7B. T cell proliferation was quantified as the dilution of eFlour450 dye and gated based on T cells cultured alone without CD3/CD28 stimulation for the undivided.

### Histology

Formalin-fixed paraffin-embedded (FFPE) lung tissue blocks were sectioned at 4-μm thickness by the Tissue Core at Moffitt Cancer Center. Hematoxylin and eosin (H&E)-stained sections were prepared by the Tissue Core immediately after sectioning. All the H&E-stained slides were imaged on an Aperio™ ScanScope AT2 (Leica Biosystems) and analyzed using GLASS-AI^19^.

### Mass spectrometry reagents and chemicals

All solvents and chemicals are LC-MS grade. Ammonium hydroxide, urea, HEPES, ammonium acetate, ammonium carbonate, ammonium formate, sodium orthovanadate, sodium pyrophosphate tetrabasic decahydrate, β-glycerophosphate disodium salt hydrate, and formic acid were obtained from Sigma Aldrich (St. Louis, MO). Water, isopropyl alcohol (IPA), methanol (MeOH), and acetonitrile (ACN) were obtained from VWR (Radnor, PA). Trifluoroacetic acid (TFA) was obtained from Fisher Scientific (Waltham, MA). The Metabolomics Quality Control (QC) kit, containing 14 isotopically-labeled metabolite standards, was obtained from Cambridge Isotope Laboratories (Tewksbury, MA) and used as an internal standard (IS). For lipidomic analysis, the Splash Lipidomix Mass Spec Standard (Avanti Polar Lipids, Alabaster, AL) kit was used as the IS.

### Multi ‘Omic Sample Preparation of Lung Tissue

Frozen tissue samples were pulverized and maintained on ice for the duration of the liquid-liquid extraction. Pulverized tissues had 2 µL of the metabolomics IS immediately followed by 500 µL of cold 80% MeOH, previously chilled for 1 hour at -80° C, for protein precipitation. Samples were vortexed thoroughly and incubated at -20° C for 30 minutes before centrifugation at 18,800 × g (Microfuge 22R, Beckman Coulter) at 4° C for 10 minutes. After centrifugation, the supernatant was transferred to a new microcentrifuge tube and dried by vacuum concentration at room temperature (SpeedVac, Thermo). Dried samples were stored in -80° C pending analysis.

The protein pellets were further processed for lipid analysis using 3 µL of the lipid IS and extracted using 300 µL of 100% IPA and vortexed thoroughly for 30 seconds. Extracts were incubated for 30 minutes at -20° C before centrifugation at 18,800 × g at 4° C for 10 minutes. The supernatant was transferred to a new microcentrifuge tube after centrifugation and dried under vacuum concentration at room temperature. Dried lipid samples were stored in -80° C pending analysis.

Untargeted metabolomic analysis was performed on UHPLC-HRMS using a Thermo Scientific Vanquish interfaced with a Q Exactive HF high-resolution mass spectrometer (Thermo Scientific, San Jose, CA). Chromatographic separation was on an Atlantis Premier BEH Z-HILIC VanGuard FIT column (2.1 mm ID x 150 mm length, 2.5 µm particle size) (Waters, Milford, MA). Gradient conditions were adapted from previously published methods^50^. Briefly, mobile phase A was aqueous 10 mM ammonium carbonate with 0.05% ammonium hydroxide, and mobile phase B was ACN. The run began at 80% B, with a linear decrease over 13 minutes to reach 20% B. It then held at 20% B for 2 minutes, before returning to the original conditions within 0.1 minutes, where it equilibrated for an additional 4.9 minutes, bringing the total run time to 20 minutes. The column was maintained at a temperature of 30° C with a flow rate of 250 µL/min. All samples were acquired in positive and negative ionization mode with a 2 µL injection volume for each analysis. Mass spectrometry was performed using heated electrospray ionization (HESI) operating at a spray voltage of 3.5 kV and 3.0 kV for positive and negative mode, a capillary temperature of 325° C, sheath gas and auxiliary gas of 50 and 10 arbitrary units (au), respectively. The mass range for acquisition was 65 – 900 m/z in positive and negative ionization mode with a mass resolution of 120,000.

Lipidomic acquisition was performed using UHPLC with tandem MS (LC-MS/MS) on a Q Exactive HF coupled with a Vanquish UHPLC. Chromatographic separation was conducted on Brownlee SPP C18 column (2.1 mm x 75mm, 2.7 µm particle size, Perkin Elmer, Waltham, MA) using mobile phase A, which was H_2_O containing 0.1% formic acid and 1% of 1M ammonium acetate and mobile phase B, which was 1:1 ACN: IPA with 0.1% formic acid and 1% of 1M ammonium acetate. The gradient separation began with column equilibration for 2 min at 35% B then a linear increase for 6 min to 80% B. A second linear increase for 14 min to 99% B was performed and held isocratic for 14.1 min B before returning to equilibrium at 35% B for 3.9 min. The flow rate was maintained at 0.400 mL/min. The lipid profile was monitored using data dependent tandem mass spectrometry for the 5 highest intensity ion signals (top 10) in positive and negative mode in separate experiments. Mass spectrometry was performed with HESI operating with a sheath gas of 50 au, auxiliary gas 10 of au, and spray voltage of 3.5 kV. The capillary temperature was maintained at 325° C. The mass range for the acquisition was 120-1000 *m/z* with a mass resolution of 120,000 for MS and 30,000 for MS/MS in positive and negative mode acquisitions. Isolation window width of 1.2 *m/z* was used for MS/MS with a stepped normalized collision energy (NCE) of 20, 30 and 40 au with a dynamic exclusion of previously sampled peaks for 8 sec.

After pulverization and metabolite/lipid extractions, the protein pellets were resuspended under sonication (Bioruptor) in denaturing aqueous lysis buffer and the proteins were extracted in denaturing aqueous lysis buffer containing 8M urea, 20 mM HEPES (pH 8), 1 mM sodium orthovanadate, 2.5 mM sodium pyrophosphate and 1 mM β-glycerophosphate. Bradford assays were carried out to determine the protein concentration for each sample. The proteins were reduced with 4.5 mM dithiothreitol (DTT) and alkylated with 10 mM iodoacetamide (IAAm) prepared as aqueous 10X stock solutions. Trypsin digestion (enzyme-to-substrate ratio 1:20) was carried out at room temperature overnight; another aliquot of trypsin is added for an additional 2 hour digestion. To quench proteolysis, tryptic peptides were then acidified with aqueous 1% trifluoroacetic acid (TFA) and desalted with C18 Sep-Pak cartridges (Waters) according to the manufacturer’s procedure.

Proteolytic peptides from each sample were labeled with TMTPro reagent (see design below). Label incorporation was checked by LC-MS/MS and spectral counting to verify that ≥95% label incorporation was achieved for each channel. The 16 samples were then pooled and lyophilized. The TMT Experimental Design used the following layout:

*Kras^G12D/+^* Ad-SPC-Cre: 126, 127C, 128C, 129C, 130C

*TAp73^Δtd/Δtd^*;*Kras^G12D/+^* Ad-SPC-Cre: 127N, 128N, 129N, 130N, 131N

*Kras^G12D/+^* Ad-CMV-Cre: 131C, 132C, 133C

*TAp73^Δtd/Δtd^*;*Kras^G12D/+^* Ad-CMV-Cre: 132N, 133N, 134N

After lyophilization, the peptides were re-dissolved in 400 µL of aqueous 20 mM ammonium formate, (pH 10.0); bRPLC A solvent was aqueous 5 mM ammonium formate, 2% acetonitrile, pH 10.0 and bRPLC B solvent was aqueous 5 mM ammonium formate, 90% acetonitrile, Ph 10.0. The basic pH reversed phase separation was performed on a BEH C18 XBridge column (4.6 mm ID x 100 mm length column packed 3.5 µm particles, 130 Å pore size, Waters). The peptides were eluted using the following gradient: 5% B for 10 minutes, 5% - 15% B in 5 minutes, 15-40% B in 47 minutes, 40-100% B in 5 minutes and 100% B held for 10 minutes, followed by re-equilibration at 1% B. The flow rate was 0.6 ml/min, and 96 fractions were collected (1 minute per well) and concatenated into 24 for protein expression or 12 for post-translational modifications. Vacuum centrifugation (Speedvac, Thermo) was used to dry the peptides.

A nanoflow ultra high performance liquid chromatograph (RSLC, Thermo, San Jose, CA) coupled to an electrospray bench top orbitrap mass spectrometer with high field asymmetric waveform ion mobility spectrometry for charge state selection (Orbitrap Exploris 480 with FAIMS, Thermo, San Jose, CA) was used for liquid chromatography tandem mass spectrometry peptide sequencing experiments. The sample was first loaded onto a pre-column (C18 PepMap100, 100 µm ID x 2 cm, 5 µm particles, 100Å pores) and washed for 8 minutes with aqueous 2% acetonitrile and 0.1% formic acid (solvent A). The trapped peptides were eluted onto the analytical column, (C18 PepMap100, 75 µm ID x 25 cm, 2 µm particles, 100 Å pores, Thermo, San Jose, CA). The 120-minute gradient was programmed as: 95% solvent A (aqueous 2% acetonitrile and 0.1% formic acid) for 8 minutes, solvent B (90% acetonitrile and 0.1% formic acid) from 5% to 38.5% in 90 minutes, then solvent B from 50% to 90% B in 7 minutes and held at 90% for 5 minutes, followed by solvent B from 90% to 5% in 1 minute and re-equilibrated at 5% B for 10 minutes. The flow rate on the analytical column was 300 nl/min. Cycle time was set at 1.5 seconds for data dependent acquisition using two FAIMS compensation voltage values (−45 V and -65 V). Spray voltage was 2,100 V, and capillary temperature was 300 °C. The resolution values for MS and MS/MS scans were set at 120,000 and 45,000 respectively. Dynamic exclusion was set to 15 seconds for previously sampled peptide peaks.

### ‘Omic Data Processing and Analysis

Metabolomics data were converted to open source mzXML files using Raw Converter (Scripps) and then imported into MZmine 3.53^51^ for feature finding and alignment. Mass detection was set to a 10 ppm window with a 0.25 min RT tolerance. Metabolite identification was performed using an in-house RT standard library. Aligned and annotated data were exported using peak height values. For each positive and negative ion mode dataset, rows with fewer than 2 high-quality pre-gap-filled features were removed as low quality. Each dataset was then normalized separately with iterative rank order normalization, IRON^47^, excluding heavy-labeled and unidentified rows from median sample identification (findmedian --pearson --iron-exclusions=unidentified_row_ids.txt --iron-spikeins=heavy_row_ids.txt) and normalization training (iron_generic --proteomics --iron-exclusions=unidentified_row_ids.txt --iron-spikeins=heavy_row_ids.txt). Normalized positive and negative ion mode datasets were then merged together and log_2_ transformed, treating zero abundances as missing data. Each row was then annotated, through fuzzy compound name matching, against our internal manually curated HMDB, KEGG, and PubChem identifier table.

Peak picking and alignment of lipidomics data were performed using LipidSearch 4.2.2.1^52^ using the HCD targeted database with a Q Exactive product search of 6 ppm for the precursor and product mass tolerances, a 1.0% MS/MS intensity threshold, and an m-Score threshold of 2. Quantitation mass tolerance of 6 ppm and a RT tolerance of ± 0.5 min was used. Alignment of the peak lists was performed using an LC-MS product search of the mean with an RT tolerance of 0.2 min and an m-Score threshold of 5.0. The aligned peak list was further filtered based on the most commonly observed adduct(s) for each lipid class. The filtered and aligned feature list was exported using peak height values and processed for normalization using the same IRON pipeline described above for metabolomics datasets.

MaxQuant (version 1.6.14.0)^53^ was used to identify peptides and quantify the TMT reporter ion intensities. Proteins were searched against murine entries in the UniProt database downloaded in April 2023. Up to 2 missed trypsin cleavage were allowed per peptide. Carbamidomethyl cysteine was set as a fixed modification, and methionine oxidation was set as a variable modification. Both peptide spectral match (PSM) and protein false discovery rate (FDR) were set at 0.01 using reverse sequences for the decoy. The match between runs feature was activated to enable identification from each LC-MS/MS run to be carried to other runs. Similar database searches were performed in Mascot (Matrix Science) to support data upload to PRIDE/ProteomeXchange.

Data analysis of all ‘omics datasets were performed using MetaboAnalyst 6.0^54^ using one factor statistical analysis. All missing values were replaced with 1/5^th^ of the limit of detection (LOD) and feature lists were further filtered using interquartile range filtering for those containing more than 5,000 features. Pathway analyses for metabolomics and lipidomics datasets was performed using relative-betweenness centrality using the KEGG database.

### Targeted Eicosanoid Metabolomics

Isotopically-labeled standards at 99% purity were purchased from Cayman Chemical (Ann Arbor, MI), unless otherwise noted, to generate a spike-in IS: Docosapentaenoic Acid-d_5_, α-Linolenic Acid-d_5_, Thromboxane B2-d_4_, Arachidonic Acid-d_5_, Prostaglandin D2-d_4_, (±)11(12)-EET-d_11_, and 20-HETE-d_6_. IS mixture was generated in UHPLC grade MeOH for an in-sample spiked concentration of 5 μg/mL. Fresh IS mix was prepared prior to each sample preparation and stored at -80° C until needed.

Homogenization of frozen murine tumors was performed as described above and isolated macrophages were homogenized via sonication in preparation for extraction. Human serum was aliquoted at 150 μL per sample for extraction. Sample extraction for all biological matrices followed the same metabolomics protocol described above adjusted to maintain sample to extraction volume ratio. Metabolite extracts from tissue and cells were reconstituted with 80% MeOH at a protein concentration of 100 μg in 20 uL volume. Metabolite extracts from serum were reconstituted at 20 μL in 80% MeOH.

Targeted analysis was performed on UHPLC using a Thermo Scientific Vanquish interfaced with a Q Exactive HF high-resolution mass spectrometer using a modified version of a previously published LC-MS/MS method^23^. Chromatographic separation was on performed by injecting 5 µL of reconstituted samples onto a Waters ACQUITY UPLC BEH C18 column (2.1 mm ID x 150 mm length, 1.7 µm particle size, 130 Å pore size). Gradient conditions were set as follows: mobile phase A was 0.005% acetic acid and 5% ACN in water, and mobile phase B was 0.005% acetic acid in ACN. Chromatography began at 35% B, where it remained isocratic for 1 minute before a linear increase over 9 minutes to 98% B. It remained isocratic at 98% B for 0.5 minutes, before returning to the starting conditions within 0.1 minutes, where it equilibrated for an additional 0.4 minutes, with a total run time of 12 minutes. The column temperature was maintained at 55 °C with a flow rate of 300 µL/min. Mass spectrometry was performed in negative ion mode using HESI operating at a spray voltage of 3.0 kV, a capillary temperature of 320° C, sheath gas and auxiliary gas of 50 and 10 arbitrary units (au), respectively. The mass spectrometer operated in full scan and parallel reaction monitoring (PRM) modes. Full scan operated at a mass range of 100 – 600 *m/z* with a mass resolution of 120,000 and PRM mode was acquired using a mass resolution of 30,000 and a normalized collision energy (NCE) of 20. Metabolite parameters are described in Table S6.

Data were processed using Thermo Scientific XCalibur 3.0 processing software for feature detection, peak area integration, and feature list generation. Peak integration was performed using a 30 second chromatographic isolation window with a mass tolerance of 10 ppm using Genesis peak detection with five-point smoothing. Peak integration was obtained using the targeted precursor ion and the MS/MS product ions for higher selectivity. Statistical analyses were performed using the same metabolomics workflow described above.

### Ion Mobility Mass Spectrometry Imaging

Intact lung lobes from each genotype initiated with each viral vector were prepared for ion mobility-matrix assisted laser/desorption ionization (MALDI) mass spectrometry imaging (IM-MALDI-MSI) using the protocol as previously published^55^. Briefly, murine tissue lobes were insufflated with Milestone Cryoembedding Compound (MCC)(Milestone Medical, 6705MILE01) and immediately snap frozen in liquid nitrogen (LN_2_). Frozen lung tissues were cryosectioned at 5 μm thickness and mounted either on standard microscopy slides for H&E staining and immunofluorescence or on indium-tin oxide (ITO) coated slides for IM-MALDI-MSI. ITO slides were sprayed with N-(1-naphthyl)ethylenediamine Dihydrochloride (NEDC)(Tokyo Chemical Industry, N0869) at 7 mg/mL in aqueous 50% MeOH using an HTX M5 Sprayer before being acquired on a Bruker timsTOF fleX MALDI ion mobility mass spectrometer at 20 μm imaging across the entire lung sections. Imaging data were processed using SCiLS Lab (v. 12.01 Core, Bruker) by normalizing to root mean square and co-registering H&E-stained images to identify tumor regions of interest.

### Immunofluorescence

Slides of frozen lung tissue serially sectioned with the sections analyzed by MALDI-MSI were allowed to warm to room temperature for 30 minutes. The slides were simultaneously fixed and dehydrated by incubating in Signature Series Pen-Fix (Epredia 22-046340) for 15 minutes and then rinsed in PBS. Antigen unmasking solution (Vector Laboratories H-3300) was prepared according to the manufacturer’s directions, supplemented with 0.05% Tween 20 (Fisher BP337), and placed in a steamer to warm up. The fixed slides were then incubated in the pre-warmed antigen retrieval solution in the steamer for 22 minutes, allowed to cool at room temperature for 20 minutes, and rinsed in PBS. Slides were blocked in PBS plus 5% goat serum (Vector Laboratories S-1000-20) and 0.2% Triton X-100 (Sigma-Aldrich X100) for one hour at room temperature and then incubated with rabbit anti-CD45 (Cell Signaling 70257, RRID:AB_2799780) diluted 1:50 in blocking buffer overnight at 4° C. Slides were rinsed in PBS, incubated with goat anti-rabbit AF488 (Invitrogen A-11008, RRID:AB_143165) diluted 1:500 in blocking buffer containing 1 µg/mL DAPI (Thermo Scientific 62247) for one hour at room temperature, and rinsed with PBS again before coverslipping with ProLong Diamond Antifade Mountant (Molecular Probes P36961). Whole-slide images were acquired using the PhenoCycler-Fusion system (Akoya Biosciences).

### Immunohistochemistry

With the help of the Tissue Core at Moffitt Cancer Center, 4-μm sections were cut from FFPE tissue blocks and were stained using the Ventana Discovery XT automated system and associated proprietary reagents (Ventana Medical Systems) per the manufacturer’s protocol. The slides were stained with rabbit anti-CD3 (Spring Bioscience M3074, RRID:AB_1660772) and rabbit anti-CD11b, (LSBio LS-C141892, RRID:AB_10947510). Immunostained slides were counterstained with hematoxylin, dehydrated, and mounted following standard protocols.

### Analysis of immunostained slides

Immunostained slides were scanned using the Aperio™ ScanScope AT2 (Leica Biosystems) with a 20x/0.75NA objective lens. Images were stored in Aperio’s Eslide manager software in SVS file format. The SVS whole slide images were imported into Definiens Tissue Studio v4.2 suite for tumor segmentation and analysis. In Tissue Studio, a machine learning algorithm (Composer) was used to segment each tissue section image into Tumor and Non-Tumor. This segmentation was further refined using a minimum size feature and manual corrections by an experienced image analyst. Next, a cell detection algorithm was used to identify all cells within the Tumor compartment and classify them as positive or negative for the marker of interest based on staining intensity. The cell count data for each image was exported into Microsoft Excel where the density of immunostained cells was calculated as the number of positive cells divided by the total tumor area identified per section.

### scRNA-seq sample preparation

After perfusing the heart and lungs with PBS, fluorescent tumors were resected and kept on ice until ready for processing. To generate a single-cell suspension, the tumors were first minced with a razor blade and then digested for 15 minutes at 37° with 2 mg/mL Collagenase I (Gibco 17100-017), 2 mg/ml Dispase II (Roche 04942078001), and 25 U/mL DNase I (Sigma D4527) diluted in Eagle’s Minimum Essential Medium (VWR 10128-610). The digestion media was neutralized by 3- to 4-fold dilution with PBS containing 2% fetal bovine serum (FBS) and filtered through 40 µm mesh. The red blood cells were lysed (Biolegend 420301) and the remaining cells were resuspended in PBS with 0.4 mg/mL BSA (Promega R3961) to a final concentration of 500-1000 cells/µl.

### scRNA-seq encapsulation and library prep

Single-cell RNA-sequencing was performed using the 10X Genomics Chromium system. A single-cell suspension derived from dissociated mouse tumor tissue was analyzed for viability using the Nexcelom Cellometer K2 and then loaded onto the 10X Genomics Chromium Single Cell Controller at a concentration of one thousand cells per microliter in order to encapsulate approximately 3,000 to 6,000 cells per sample. Briefly, the single cells, reagents, and 10x Genomics gel beads were encapsulated into individual nanoliter-sized Gel beads in Emulsion (GEMs) and then reverse transcription of poly-adenylated mRNA was performed inside each droplet. The cDNA libraries were then completed in a single bulk reaction using the Chromium Single Cell Gene Expression 3’ v3 Reagent Kit, (10X Genomics Chromium NextGEM Single Cell 3’ Reagent Kit v3.1) and approximately 40,000 sequencing reads per cell were generated on the Illumina NextSeq 500 instrument (Illumina NovaSeq 6000 instrument).

### scRNA-seq data processing, filtering, batch effect correction, and clustering

A customized reference genome was created using the GRCm38 mouse transcriptome with the *mkref* module of Cell Ranger (v6.0, 10X Genomics). Raw sequencing reads from single cells were aligned to this reference and processed with the count module of Cell Ranger. Gene-barcode matrices, including only barcodes with UMI counts meeting quality thresholds, were imported into Seurat for downstream analysis. Genes detected in fewer than three cells were excluded, as were cells with fewer than 200 detected genes or more than 10% mitochondrial UMIs. Potential doublets were identified using Scrublet, DoubletFinder, scDblFinder, and doubletCells implemented in scran, assuming a doublet rate of 0.08% per 1,000 sequenced cells. Cells flagged as doublets by at least two methods were removed. Raw UMI counts were log-normalized, and the top 5,000 variable genes were identified using the “vst” method in the *FindVariableFeatures* function of Seurat. To avoid clustering based on V(D)J transcripts, T-cell receptor and immunoglobulin genes were excluded from the variable gene list. Cell cycle phase scores (S and G2/M) were calculated using the *CellCycleScoring* function.

To address batch effects, individual samples were integrated using the *FindIntegrationAnchors* and *IntegrateData* functions in Seurat. Integration used 8,000 anchor genes and 40 dimensions of canonical correlation analysis (CCA). Dimensional reduction was performed with diagonalized CCA, and L2-normalization was applied to canonical correlation vectors to project all samples into a shared space. Mutual nearest neighbors (MNNs) between datasets served as “anchors” to encode cellular relationships and compute correction vectors for integration.

The integrated data were scaled using the *ScaleData* function, regressing out total UMI counts, mitochondrial UMI percentages, and cell cycle scores (S and G2/M). A shared nearest neighbor (SNN) graph was constructed using the top 40 principal components, and clusters were identified with the Louvain algorithm (*FindCluster*) at a resolution of 0.6, yielding 38 clusters. For visualization, Uniform Manifold Approximation and Projection (UMAP) was generated using the RunUMAP function.

### Mouse cluster annotation

Cluster-specific differential expression analysis was performed using the *FindAllMarkers* function in Seurat with default parameters. Genes with a log2(fold-change) > 0.25 and a Bonferroni-corrected p-value < 0.05 were considered differentially expressed. Cluster annotations were based on comparisons of cluster-specific genes to canonical markers for the following major cell types: macrophage (*C1qc*, *Mafb*, *Cd68*), B cell (*Pax5*, *Cd79a*, *Cd19*), epithelial (*Krt6a*, *Dsp*, *Krt18*), T cell (*Cd3e*, *Cd3d*, *Cd4*, *Cd8a*), endothelial (*Pecam1*, *Cdh5*, *Tie1*), monocyte (*Ly6c2*, *Nr4a1*, *Fcgr1*, *Lyz2*), NK cell (*Klrb1c*, *Ncr1*, *Eomes*), fibroblast (*Col1a1*, *Col1a2*, *Col5a1*), mregDC (*Fscn1*, *Ccr7*), pDC/MonoDC (*Siglech*, *Bst2*, *Tcf4*, *Cd209a*), neutrophil (*S100a8*, *Mmp9*, *Cxcr2*), cDC1 (*Xcr1*, *Cadm1*, *Itgae*). Expression of marker genes was visualized using UMAP plots and violin plots based on log-normalized UMI counts. The 38 clusters were further grouped into three major categories: T/NK cells, myeloid cells, and lung epithelial cells.

### Deconvolution of bulk RNAseq

We had previously performed RNA sequencing on total RNA from snap-frozen tumors isolated from *Kras^G12D/+^*;*Rosa^mG/mG^*and *TAp73^Δtd/Δtd^*;*Kras^G12D/+^* mice infected with Ad-CMV-Cre (3 and 4 tumors, respectively; available at GEOXXX). Complementary DNA libraries were prepared using the Clontech Smarter Stranded V2 kit, and over 100 million paired-end reads were generated per sample using v2 chemistry on an Illumina NextSeq 500 sequencer. The RNAseq reads were aligned onto the mouse genome build UCSC mm10 (NCBI 38) using the STAR aligner^56^. Gene expression was quantified using RSEM^57^, normalized for library size and transformed using voom^58^, and analyzed by limma^59^ to determine differentially expressed genes according to linear models. We performed Gene Set Enrichment Analysis of this data set using signatures taken from the 12 major cell types identified in the single-cell sequencing data (p-adj ≤ 0.05, fold change > 1.5; Table S7).

### Pathway analysis

Enrichment of pathways among differentially expressed proteins (FC>1.5, p<0.05; Fig. 3D-3E) was carried out with mSigDB^60,61^ by mapping the mouse gene identifiers to the human orthologs and computing the overlaps with the human hallmark gene sets or canonical pathways with a FDR q-value < 0.05. Enrichment of pathways among differentially expressed transcripts identified from single-cell sequencing (log2FC>0.25, p-adj<0.05; Fig. 5E-5F) was carried out with Enrichr^62^ by computing the overlaps with the WikiPathways 2024 Mouse gene-set library^63^ with an p-adj<0.05. For characterizing different alveolar macrophage subtypes identified by single-cell sequencing (Supplementary Fig. S6E), the single sample GSEA algorithm^61,64^ was applied to the rank-normalized list of genes expressed in at least 10% of the cells of a given subtype.

### T cell Isolation and Macrophage Co-Culture

Spleens from wild-type mice were harvested and CD3^+^ T cells were negatively enriched according to the manufacturer (Magnisort Mouse CD3 T cell Enrichment Kit, ThermoFisher 8804-6820-74). CD3^+^ T cells were stained with 10 μM eBioscience Cell Proliferation Dye eFluor450 (Invitrogen 65-0842-85) at 4E6 cells/ml for 15 min in a 37°C water bath followed by a PBS wash to remove excess dye and quenched on ice for 10 min. Labeled eF450^+^ CD3^+^ T cells were counted and plated in co-cultures. Co-cultures were maintained in 200 μl of total media in round bottom tissue-culture treated 96-well plates. For all co-cultures, 1E5 T cells/well were added and 1E5 Siglec-F^+^ macrophages/well (isolated as above) were added and serially diluted by half for ratios of T cells:macrophages ranging from 1:1 to 1:0.125. For controls, CD3^+^ T cells were cultured alone without macrophages in the absence (negative control) or presence (positive control) of 1 μg/ml of plate bound antibodies to CD3 (BD 553058, RRID:AB_394591) and CD28 (BD 553294, RRID:AB_394763). Co-cultures were maintained at 37 °C in a humidified incubator with 5% carbon dioxide for 72 hours before being analyzed by flow cytometry as described above.

### ELISA

Interferon gamma levels in the supernatant from the co-cultured macrophages and T cells was analyzed by ELISA according to the manufacturer’s protocol (Invitrogen KMC4021). Prior to analysis, the supernatant was centrifuged at 500xg for 5 minutes at 4 °C to remove any particulate. The samples were diluted either 1:4 or 1:50 for the concentration to fall within the range of the standard curve.

### Human scRNA-seq analysis

NSCLC 10x scRNA-seq data (GSE207422^33^) was downloaded from GEO database. scRNA-seq FASTQ files were processed using the BD Rhapsody Whole Transcriptome Analysis (WTA) Pipeline to get a unique molecular identifier (UMI) matrix for each sample. Gene-barcode matrices, including only barcodes with UMI counts meeting quality thresholds, were imported into Seurat V5 for downstream analysis. Genes detected in fewer than three cells were excluded, as were cells with fewer than 200 detected genes or more than 10% mitochondrial UMIs. Raw UMI counts were log-normalized, and the top 5,000 variable genes were identified using the “vst” method in the *FindVariableFeatures* function of Seurat. To avoid clustering based on V(D)J transcripts, T-cell receptor and immunoglobulin genes were excluded from the variable gene list. Cell cycle phase scores (S and G2/M) were calculated using the *CellCycleScoring* function.

To address batch effects, individual samples were integrated using the *HarmonyIntegration* method in Seurat. The integrated data were scaled using the *ScaleData* function, regressing out total UMI counts, mitochondrial UMI percentages, and cell cycle scores (S and G2/M). A shared nearest neighbor (SNN) graph was constructed using the top 40 principal components, and clusters were identified with the Louvain algorithm (*FindCluster*) at a resolution of 1.0. For visualization, Uniform Manifold Approximation and Projection (UMAP) was generated using the RunUMAP function.

Cluster annotations were determined by comparing cluster-specific genes with canonical markers, particularly for macrophages and epithelial cell types. Expression pattern and differential expression of selected marker genes (ANXA2, S100A10 and ELOVL5) between conditions were visualized using UMAP and violin plots based on log-normalized UMI counts.

For another GEO dataset (GSE154826^32^), we used the processed 10x scRNA-seq data and its accompanying annotation to perform expression patterns and differential expression, following the same approach of downstream analyses.

### FLAG immunoprecipitation

To test the ability to immunoprecipitate FLAG-tagged TAp73, THP-1 cells were lysed with IP buffer (Pierce 87787) with protease inhibitors (Neta Scientific 11697498001). The lysates were cleared with Dynabeads Protein G magnetic beads (Fisher Scientific 10-007-D) and immunoprecipitated with Anti-FLAG M2 Magnetic Beads (Sigma M8823) after setting aside 5% of the input to analyze by Western blot. Lysates were electrophoresed on a 10% SDS PAGE gel and transferred to a nitrocellulose membrane (BioRad 1620115). Blots were probed with the anti-p73 antibody EP436Y (Abcam ab40658, RRID:AB_776999, 1:500) at 4 °C overnight, followed by incubation for 1 hr at room temperature with goat anti-rabbit HRP (Jackson ImmunoResearch 111-035-003, RRID: AB_2313567, 1:5,000). ECL Prime reagent (Cytiva RPN2232) was used for chemiluminescent detection on the Odyssey Fc imaging system (LiCOR) and analyzed using the Image Studio software (LiCOR).

### Chromatin immunoprecipitation (ChIP)

Cellular proteins were crosslinked to DNA using 1% formaldehyde and chromatin was prepared as previously described^65^. Each ChIP was performed in triplicate using either Anti-FLAG M2 Magnetic Beads (Sigma M8823) or 10 µg mouse IgG purified from mouse serum (Santa Cruz sc-2025) as a negative control for the immunoprecipitation. The recruitment of TAp73 was analyzed by qRT-PCR comparing regions with a predicted p73 binding site (described in Table S8) to nearby non-specific regions using the primers listed in Table S9.

## Data availability

New RNAseq and single-cell RNAseq data have been deposited at GEO and are publicly available as of the date of publication (GSE309911 and GSE309973, respectively). The mass spectrometry proteomics data have been deposited to the ProteomeXchange Consortium via the PRIDE^66^ partner repository with the dataset identifier PXD069509 and 10.6019/PXD069509. This study also includes analyses of existing, publicly available data, and these accession numbers are listed in the manuscript.

## AUTHORS’ CONTRIBUTIONS

**H.D. Ackerman:** Conceptualization, data curation, formal analysis, investigation, methodology, visualization, writing–original draft, writing–review and editing. **V.Y. Rubio:** Data curation, formal analysis, investigation, methodology, visualization, writing–review and editing. **A.J. Davis:** Formal analysis, investigation, visualization, writing–review and editing. **J.H. Lockhart:** Formal analysis, investigation, visualization, writing–review and editing. **N. Hackel:** Investigation. **R.V. Jimenez:** Formal analysis, investigation, writing–review and editing. **R.A. Sierra-Mondragon:** Investigation. **J.R. Baldwin:** Investigation. **C. Carr:** Investigation. **M. Reiser:** Investigation. **M. Napoli:** Investigation, visualization, writing–review and editing. **X. Yu:** Formal analysis, visualization. **C.-H. Cheng:** Formal analysis, visualization. **P.A. Stewart:** Formal analysis. **S. Acevedo-Acevedo:** Resources, writing–review and editing. **R. Checker:** Investigation, writing–review and editing. **I. Grammatikakis:** Investigation, writing–review and editing. **X. Su:** Investigation. **Y. Wu:** Investigation. **T. Gould:** Investigation. **A. Bailey:** Resources. **L.A. Robinson:** Resources, writing–review and editing. **E.B. Haura:** Resources. **J.M. Koomen:** Supervision, writing–review and editing. **P.C. Rodriguez:** Supervision, writing–review and editing. **E.R. Flores:** Conceptualization, funding acquisition, supervision, writing–review and editing.

## ACKNOWLEDGEMENTS

Initial funding for this work was provided by an NIH/NCI R35CA197452 to E.R.F. and the Miles for Moffitt Foundation. Current and ongoing support for this work has been provided by NIH/NCI P01CA250984 to E.R.F. Mass spectrometry imaging has been supported by NIH instrumentation funding and matching institutional funds (S10OD032419 to J.M.K.). The funders played no role in the study design, data collection, analysis, and interpretation of data, or the writing of this manuscript. J.H.L. was supported by the Integrated Program in Cancer and Data Science postdoctoral training fellowship (T32CA233399) from the National Cancer Institute. We thank Drs. Eric Haura, Lary Robinson, and Gina DeNicola for providing access to human data or samples. This work has been supported in part by the Analytic Microscopy Core, the Biostatistics and Bioinformatics Shared Resource, the Comparative Medicine Core, the Flow Cytometry Core, the Molecular Genomics Core, the Proteomics and Metabolomics Core, the Quantitative Imaging Core, the Small Animal Imaging Core, and the Tissue Core at the H. Lee Moffitt Cancer Center & Research Institute, an NCI-designated Comprehensive Cancer Center (NCI P30CA076292). The contents of this work are solely the responsibility of the authors and do not necessarily represent the official views of the NCI or NIH.

## SUPPLEMENTARY DATA

Document S1. Supplementary Figures S1–S8, Supplentary Tables S8 and S9

Table S1. Eicosanoid-targeted metabolomics from mouse tumors, related to Figure 3

Table S2. Eicosanoid-targeted metabolomics from alveolar macrophages, related to Figure 6

Table S3. Eicosanoid-targeted metabolomics from THP-1 cells, related to Figure 7

Table S4. Eicosanoid-targeted metabolomics from NSCLC patient serum, related to Figure 7

Table S5. Southern blot probe sequences, related to Figure 1

Table S6. Metabolite parameters for eicosanoid-targeted assay, related to Figures 3, 6, and 7

Table S7. Gene signatures for major cell types from single-cell sequencing, related to Supplementary Figure 6

## Notes

### Competing Interest Statement

The authors have declared no competing interest.

