## Supplementary Figures S1-S8, Tables S8 and S9 for "TAp73 mediates anti-tumor immunity through regulation of lipid metabolism in the lung tumor microenvironment"

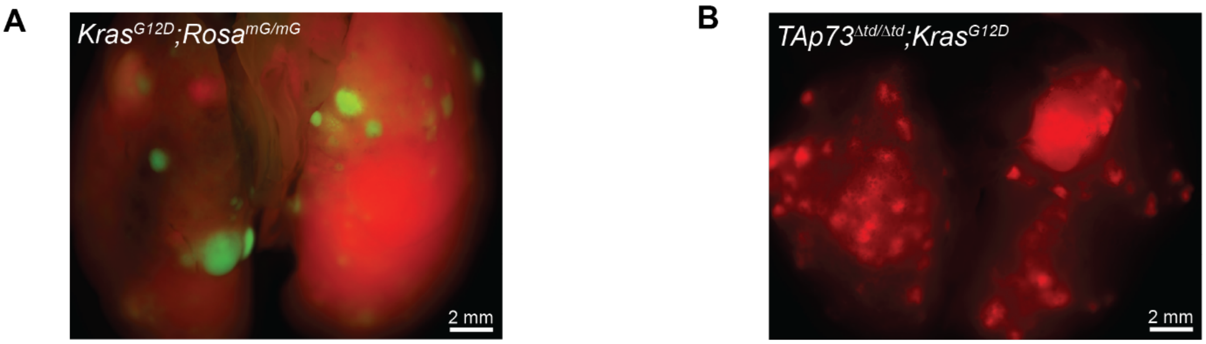


**Figure S1.** Fluorescent images of tumor-bearing lungs, related to Figure 1. **A,** Representative image of lungs from a *Kras*$^{G12D/+}$;*Rosa*$^{mG/mG}$ mouse 30 weeks after intratracheal infection with Ad-CMV-Cre. Tumors are EGFP$^{+}$ (green) while normal lung is tdTomato$^{+}$ (red) from the Cre-dependent Rosa reporter. **B,** Representative image of lungs from a *TAp73*$^{\Delta td/\Delta td}$;*Kras*$^{G12D/+}$ mouse 30 weeks after intratracheal infection with Ad-CMV-Cre. Tumors are tdTomato$^{+}$ (red) indicating Cre-dependent ablation of *TAp73*.

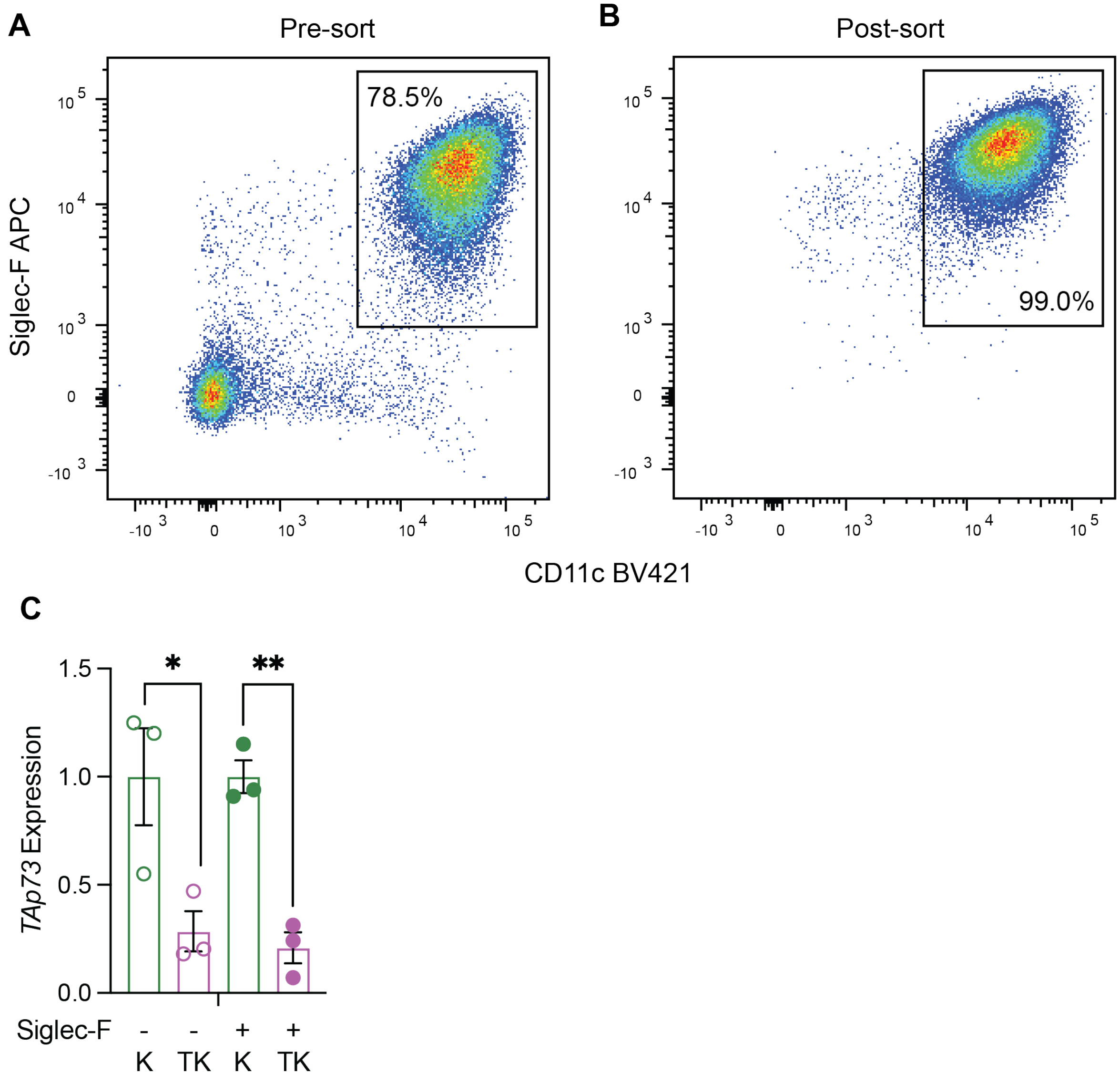


**Figure S2.** Validation of Siglec-F$^+$ macrophage isolation and *TAp73* ablation, related to Figure 2. **A-B,** Representative flow cytometry scatter plots of Siglec-F$^+$ CD11c$^+$ alveolar macrophages in the lungs of a tumor-bearing *TAp73*$^{\Delta td/\Delta td}$;*Kras*$^{G12D/+}$ (TK) mouse infected with Ad-CMV-Cre before (**A**) and after (**B**) immunomagnetic positive selection for Siglec-F$^+$ cells. The percentage is calculated based on the CD45$^+$ parent population of live single cells. **C,** qPCR of relative *TAp73* expression in Siglec-F$^+$ alveolar macrophages isolated from *Kras*$^{G12D/+}$ (K) and TK mice infected with Ad-CMV-Cre and in the remaining macrophage-depleted (Siglec-F$^-$) single-cell suspensions. Data are represented as mean ± SEM. *$p < 0.05$, **$p < 0.01$ as determined by unpaired t test.

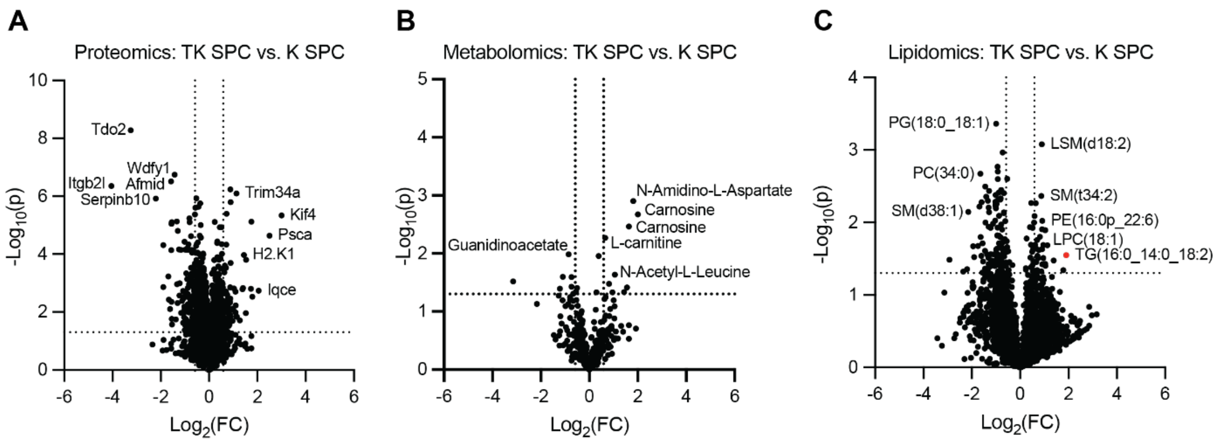


**Figure S3.** Differentially abundant features in tumors with and without tumor-specific *TAp73* ablation, related to Figure 3. **A-C,** Volcano plots representing the proteins (**A**), metabolites (**B**), or lipids (**C**) that were differentially abundant among tumors isolated from $Kras^{G12D/+}$ (K) and $TAp73^{\Delta td/\Delta td}$;$Kras^{G12D/+}$ (TK) mice infected with Ad-SPC-Cre (5 tumors per group). Red dot in **C** represents upregulated lipid in the triglyceride class.

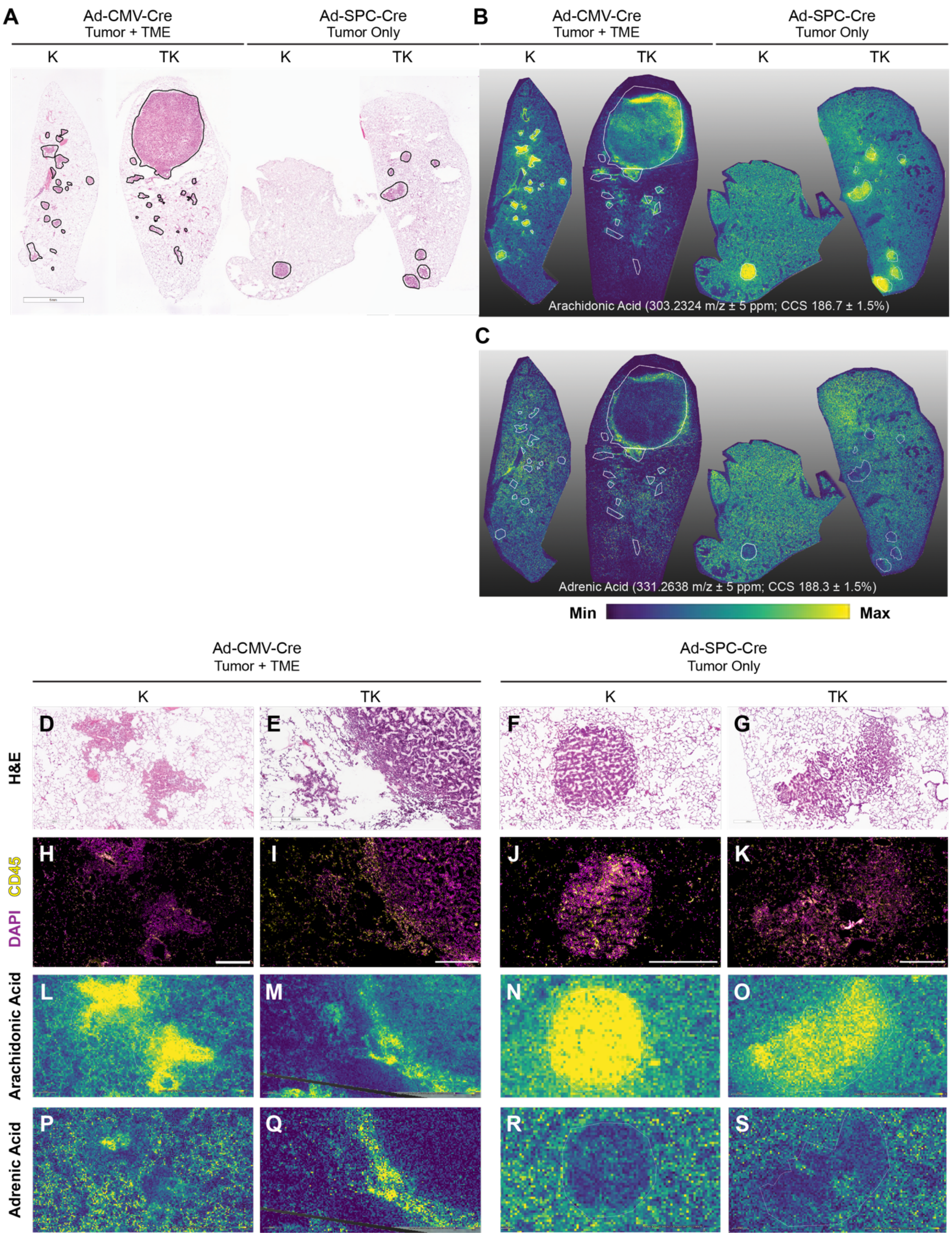


**Figure S4.** TAp73-deficient TME expresses high levels of arachidonic and adrenic acid, related to Figure 3. **A,** H&E images of the lung lobes from *Kras*$^{G12D/+}$ (K) and *TAp73*$^{\Delta td/\Delta td}$;*Kras*$^{G12D/+}$ (TK) mice infected with Ad-CMV-Cre or Ad-SPC-Cre that were analyzed by MALDI-MSI. Tumor regions are outlined in black. **B-C,** Ion images for arachidonic acid (**B**) or adrenic acid (**C**) from lung sections cut sequentially after those pictured in **A**. Areas with maximum intensity are represented in yellow while areas of minimum intensity are represented in blue. The tumor outlines from the H&E images are overlaid in white. **D-G,** Higher magnification images of regions of interest from the H&E images in **A**. **H-K,** Anti-CD45 immunofluorescence of the regions in **D-G**. Scale bars represent 500 μm. **L-O,** Arachidonic acid ion images of the regions in **D-G**. **P-S,** Adrenic acid ion images of the regions in **D-G**.

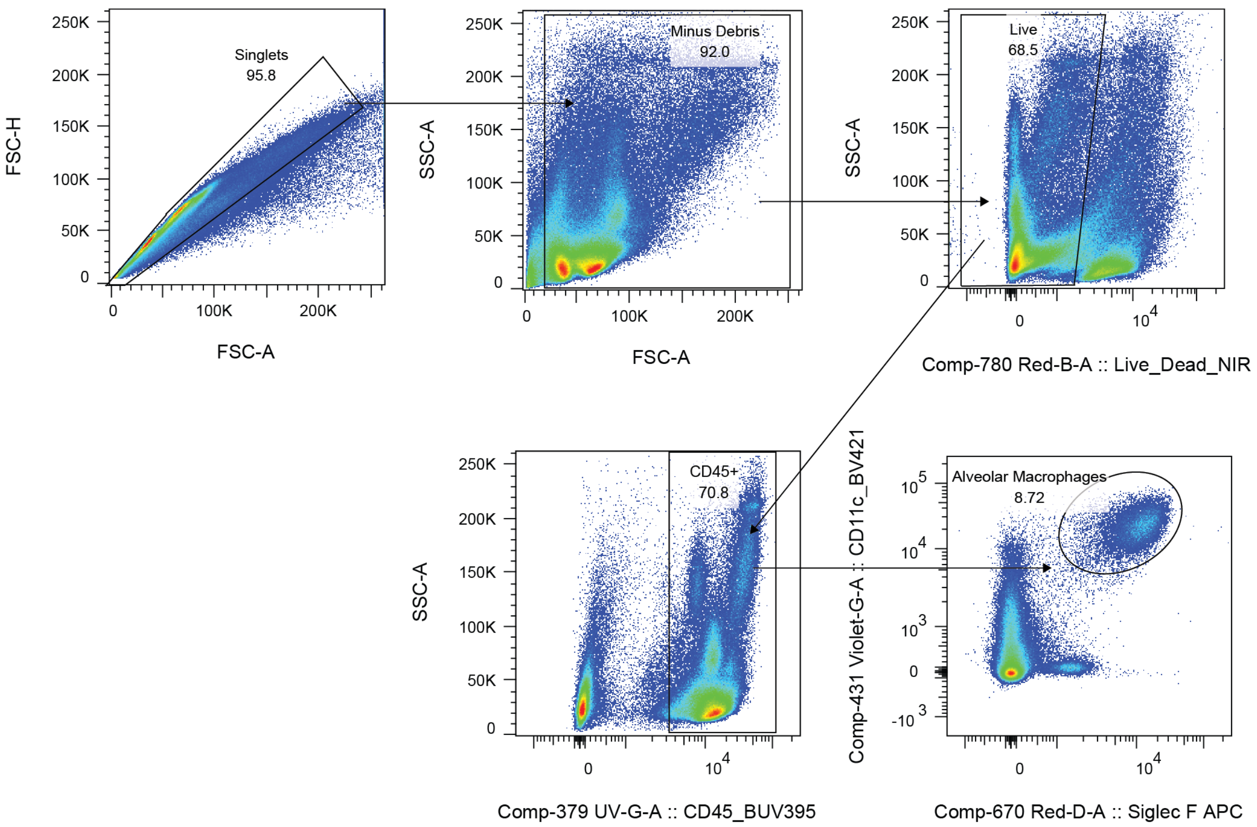


**Figure S5.** Quantification of alveolar macrophages by flow cytometry, related to Figure 4. Flow cytometric gating strategy used for quantifying Siglec-F$^+$ CD11c$^+$ alveolar macrophages.

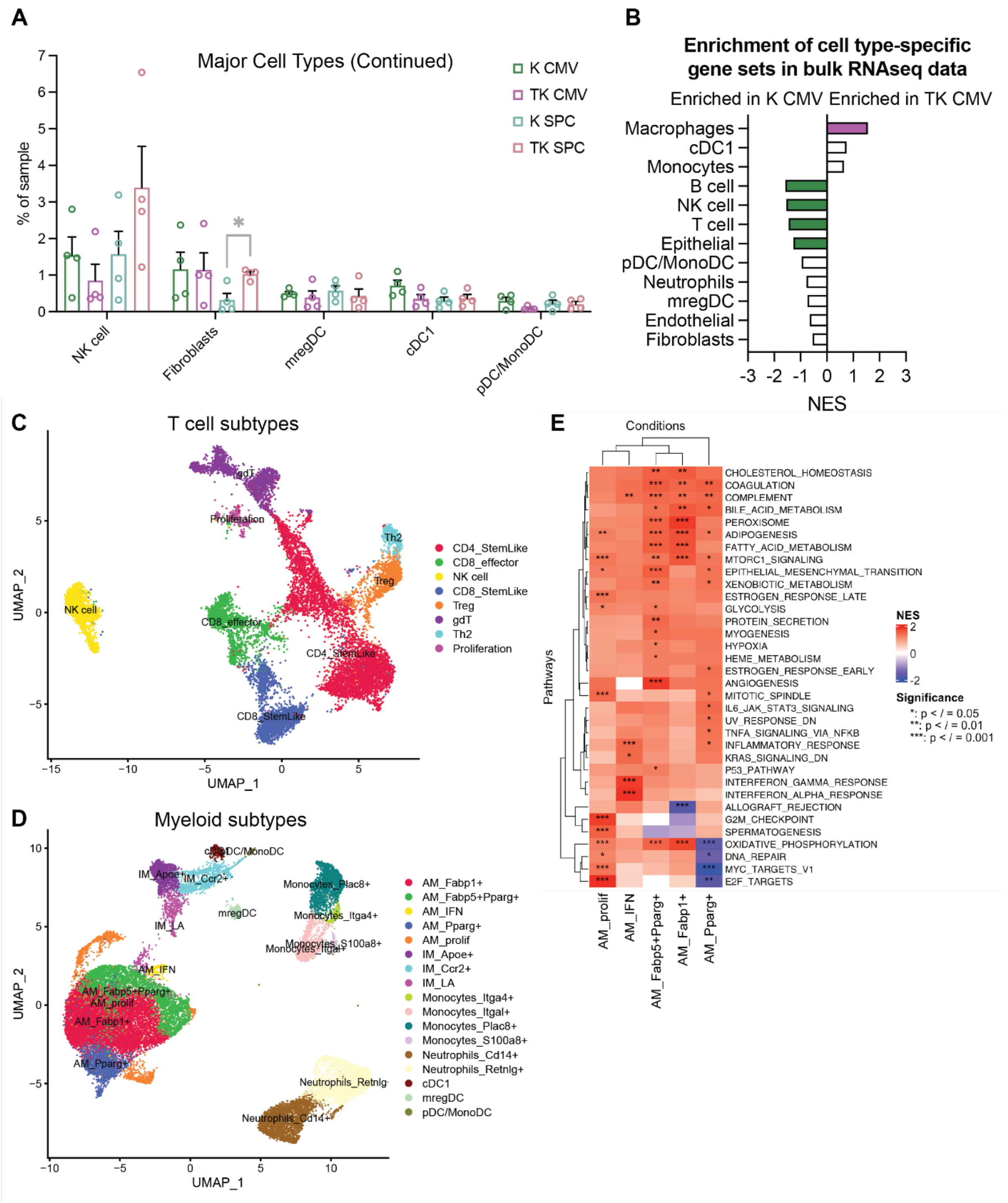


**Figure S6.** Analysis of major cell types and subtypes identified by single-cell sequencing, related to Figure 5. **A,** Continuation of Figure 5B showing the distribution of the 5 least abundant cell types within each sample across the four genotype:virus combinations. Data are represented as mean ± SEM. **p* < 0.05 by multiple unpaired t tests. **B,** Plot summarizing the normalized enrichment scores (NES) for the gene signatures derived from the 12 major cell types identified by single-cell sequencing in a separate RNA sequencing dataset from 3 K CMV and 4 TK CMV bulk tumors. The bars for gene sets that are significantly enriched (p-adj<0.05) in TK CMV are filled with purple while gene sets that are significantly enriched in K CMV are filled with green. **C,** UMAP color-coded based on 8 different T cell and NK cell subtypes. **D,** UMAP color-coded based on 17 different myeloid subtypes including macrophages, neutrophils, monocytes, and dendritic cells. **E,** Heatmap summarizing the enrichment of hallmark gene signatures across the five different subtypes of alveolar macrophages identified by single-cell sequencing. Significance is indicated by asterisks according to the adjusted p-value cutoffs.

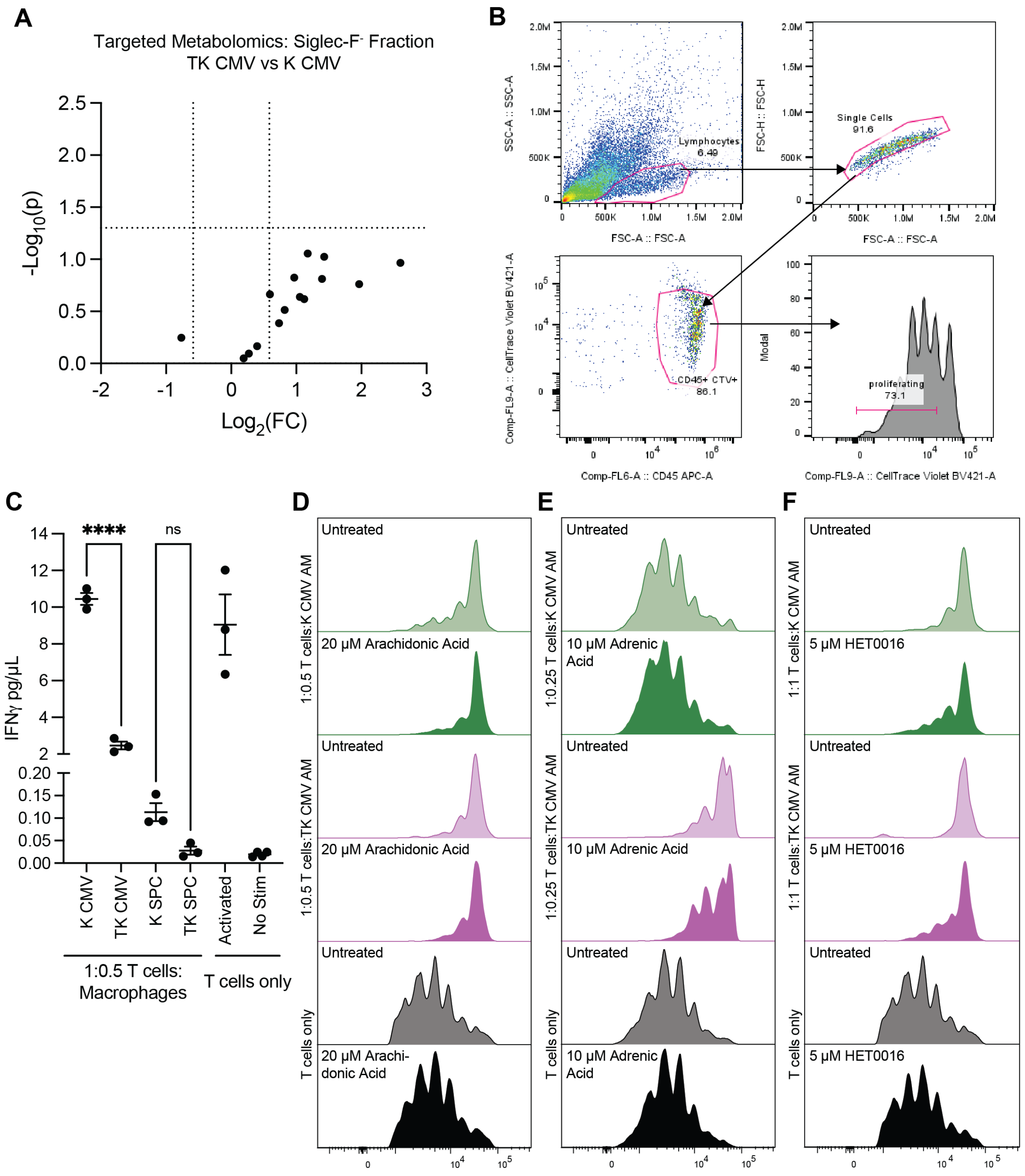


**Figure S7.** T cell activation assay, related to Figure 6. **A,** Volcano plot representing the eicosanoid-targeted metabolomics of the Siglec-F$^-$ fraction of lung cells isolated from *Kras$^{G12D/+}$* (K) and *TAp73$^{\Delta td/\Delta td}$;Kras$^{G12D/+}$* (TK) mice infected with Ad-CMV-Cre. Metabolites were considered to be significantly different if they had a fold change of at least 1.5 with $p<0.05$. **B,** Flow cytometric gating strategy used for quantifying proliferating lymphocytes. **C,** ELISA data quantifying the amount of IFNγ released by T cells after 72 hours of culture alone or in the presence of Siglec-F$^+$ alveolar macrophages isolated from tumor-bearing mice. *$p < 0.05$,

**$p < 0.01$, ***$p < 0.001$, ****$p < 0.0001$ by ANOVA followed by Tukey's multiple comparisons test. **D-F,** Representative histograms for the data in 6G-6I.

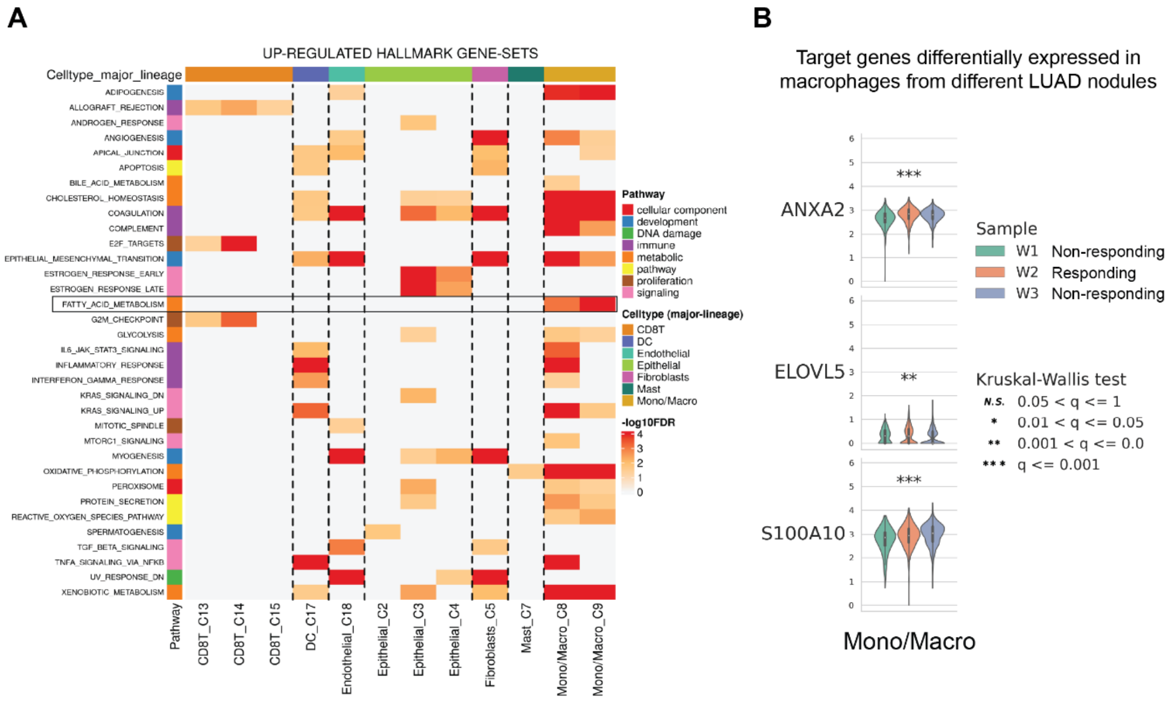


**Figure S8.** Analysis of TAp73-regulated genes and pathways in a human scRNA-seq dataset, related to Figure 7. **A,** Heatmap showing the -log$_{10}$(FDR) for hallmark gene-sets that are significantly enriched in different cell types in the scRNA-seq dataset from a LUAD patient treated with immune checkpoint inhibitor therapy (GSE146100). The figure was generated using the TISCH2 web resource (http://tisch.comp-genomics.org/). The enrichment of fatty acid metabolism in cells of the monocyte/macrophage lineage has been highlighted in the black rectangle. **B,** Violin plots showing expression levels of TAp73 target genes in macrophages from different nodules of the LUAD patient shown in (A). The figure was generated using the TISCH2 web resource.

| Element | Location | p73 binding site | | | JASPAR score |
|---|---|---|---|---|---|
| ANXA2 promoter | -5,568 to -5,552 | c**CA**t**G**Tc | ag | aA**C**At**G**t | 605 |
| S100A10 promoter | -7,629 to -7,613 | g**CA**g**G**Tc | at | gT**C**Tt**G**c | 435 |
| ELOVL5 promoter | -6,660 to -6,644 | a**CA**t**G**Ca | aa | aT**C**At**G**t | 469 |
| CYP4F3 intron 3 | 5,709 to 5,725 | c**CA**t**G**Tc | ag | gA**C**Tt**G**t | 570 |
| CYP4F2 intron 3 | 5,046 to 5,062 | c**CA**t**G**Tc | ag | gA**C**At**G**t | 636 |

**Table S8.** Predicted p73 binding sites, related to Figure 7. Table listing the locations of the p73 binding sites in the promoter or intronic regions of the indicated genes as predicted by the JASPAR Transcription Factor Binding Site Database. The locations are relative to the transcription start sites. The binding site sequences follow the display convention for the motif consensus line with mismatches indicated in red. A higher JASPAR score indicates greater confidence in the predicted binding site.

| Element | Binding Site | Primers (5'-3') |
|---|---|---|
| *ANXA2* promoter | p73 (-5,568 to -5,552) | Forward: GCACTTACATTCTGCTTGTTCTT |
| | | Reverse: TGGCCATCAATGAGGCTAAA |
| | NS (-3,188 to -3,172) | Forward: TAGGGCAAATCCAGCCTATTG |
| | | Reverse: GCCTTTCTCTTCCTTGGTCTT |
| *S100A10* promoter | p73 (-7,629 to -7,613) | Forward: GAGAGGAGAAATGCACCCTATC |
| | | Reverse: TGATCAGGCAGAACGGAATG |
| | NS (-4,850 to -4,834) | Forward: GTACCCAGGCTCTTTCCAATTA |
| | | Reverse: CACAATGGGACTGGAAGTGATA |
| *ELOVL5* promoter | p73 (-6,660 to -6,644) | Forward: ATCCCACATTCACTCTTGGC |
| | | Reverse: GTCATTCCAAGTTGTGTGATGTTT |
| | NS (-2,601 to -2,585) | Forward: CATTGTTGGATAGGTGGGTTAGA |
| | | Reverse: GATCCAATGCCAGGCTTACT |
| *CYP4F3* intron 3 | p73 (5,709 to 5,725) | Forward: CCATATTTATATTCAACCTGG |
| | | Reverse: CCTAGACATGAGACATGACAC |
| | NS (-653 to -637) | Forward: CCATTCATGGTTCCCTGGCT |
| | | Reverse: TCCAGCCCCACTGAGGATTA |
| *CYP4F2* intron 3 | p73 (5,046 to 5,062) | Forward: AGAGCACACCATCTCATGTTAG |
| | | Reverse: CCAGCACTCTCTGACATCTTC |
| | NS (-1,096 to -1,080) | Forward: TCAAGTGCCGCCAAAGT |
| | | Reverse: GCCTATCAGGGTGGATTAAAGG |

**Table S9.** Primer sequences for ChIP-qPCR, related to Figure 7. Primers were designed to amplify a short region of genomic DNA (125bp to 175bp) centered around a predicted p73 binding site or a non-specific (NS) region in the promoter or intronic region of the indicated genes. The locations are relative to the transcription start sites
